# Deciphering the mechanism underlying circRNA-mediated immune responses of western honeybees to *Nosema ceranae* infection

**DOI:** 10.1101/2020.10.25.353938

**Authors:** Huazhi Chen, Yu Du, Zhiwei Zhu, Jie Wang, Dingding Zhou, Yuanchan Fan, Haibin Jiang, Xiaoxue Fan, Cuiling Xiong, Yanzhen Zheng, Dafu Chen, Rui Guo

**Author notes:** These authors contributed equally to this work (Huazhi Chen, Yu Du).

## Abstract

*Nosema ceranae* is a widespread fungal parasite for adult honeybees, severely damaging bee health and sustainable development of apiculture. Circular RNAs (circRNAs) are a class of newly discovered noncoding RNAs (ncRNAs) that regulate a number of biological processes such as immune defense and development. In this current work, based on previously obtained whole transcriptome data, 8 199 and 8 711 circRNAs were predicted from the midguts of *Apis mellifera ligustica* workers at 7 days (AmT1) and 10 days (AmT2) post inoculation (dpi) with *N. ceranae* using bioinformatics. Additionally, in combination with transcriptome data from uninfected midguts (AmCK1 and AmCK2) (Xiong et al., 2018), 4 464 circRNAs were found to be shared by the aforementioned four groups, whereas the numbers of specifically transcribed circRNAs in each group were 1 389, 1 696, 1 019 and 1 871, respectively. Furthermore, 10 226 circRNAs were homologous to *Apis cerana cerana* circRNAs, while 20 circRNAs had homology with *Homo sapiens* circRNAs; in addition, 16 circRNAs were highly conserved in these three species. Differential expression analysis showed that 168 (306) differentially expressed circRNAs (DEcircRNAs) were identified in AmCK1 vs AmT1 (AmCK2 vs AmT2) comparison group, including 61 (143) upregulated circRNAs and 107 (163) downregulated circRNAs. Moreover, RT-qPCR results showed that the expression trend of eight DEcircRNAs was consistent with that of the transcriptome dataset. Based on GO database annotation, we observed that source genes of DEcircRNAs in AmCK1 vs AmT1 (AmCK2 vs AmT2) were engaged in 27 (35) functional terms, including two (two) cell renewal-associated terms, seven (seven) cell structure-associated terms, and one (one) immunity-associated terms. Additionally, DEcircRNA source genes in AmCK1 vs AmT1 were involved in two cell renewal-related pathways, Hippo and Wnt signaling pathways, and three carbohydrate metabolism-related pathways, galactose metabolism, starch and sucrose metabolism, fructose and mannose metabolism, only one energy metabolism-related pathway (oxidative phosphorylation pathway), three cellular immune-related pathways, endocytosis, phagosome, and lysosome, and a humoral immune-related pathway (FoxO signaling pathway). In AmCK2 vs AmT2 comparison group, more source genes of DEcircRNAs were associated with the abovementioned pathways relative to cell renewal, carbohydrate metabolism, and cellular and humoral immune pathways. In addition, 122 (234) DEcircRNAs in the host midgut at 7 dpi (10 dpi) with *N. ceranae* targeted 82 (106) miRNAs. Furthermore, 75 (103) miRNAs targeted by 86 (178) DEcircRNAs in AmCK1 vs AmT1 (AmCK2 vs AmT2) further bound to 215 (305) mRNAs. These targets could be annotated as an array of functional terms and pathways related to cellular renewal, cellular structure, carbohydrate and energy metabolism, and cellular and humoral immunity. In a word, we for the first time explored immune responses mediated by DEcircRNAs in the midguts of *A. m. ligustica* workers to *N. ceranae* infection. Our data provide a foundation for clarifying the molecular mechanism underlying immune response of western honeybee to *N. ceranae* invasion, but also a new insight into further understanding the host-pathogen interaction during bee microsporidiosis.

## 1. Introduction

Honeybees are the most important pollinators in nature, with tremendous economic and ecological value (Bromenshenk et al., 2015). *Apis mellifera ligustica*, a subspecies of *Apis mellifera*, has been commercially kept for crop pollination and production of bee products in China and many other countries (Bromenshenk et al., 2015). As a representative eusocial insect, *A. m. ligustica* is susceptible to infection by an array of pathogenic microorganisms such as bacteria, fungi and viruses. Among them, *Nosema ceranae* is an obligate intracellular fungal parasite that mainly infects the midgut tissue of adult honeybees, causing a serious chronic disease named bee microsporidiosis, which poses a huge threat to the beekeeping industry (Higes et al., 2020). Recent evidence has shown that *N. ceranae* is also invasive to western honeybee larvae (Eiri et al., 2015; BenVau et al., 2017). After being ingested by the bee host, *N. ceranae* spores germinate within the midgut, followed by swift ejection of the polar tube in the spore and penetration into the epithelium cell. The infective sporoplasm is then transferred into the host cell and begins to reproduce via exploitation of host material and energy (Panek et al., 2018; Paris et al., 2018). The spore number gradually increases and the induced pressure continues to increase. The host cells eventually burst to release the mature spores, which continue to infect neighboring epithelial cells or be expelled through feces to infect new individuals (Higes et al., 2007; Gisder et al., 2011). *N. ceranae* was first discovered in *Apis cerana* (Fries et al., 1996), after which it spread to western honeybee colonies in European countries and Taiwan Province, China (Huang et al., 2007; Chen et al., 2008). At present, *N. ceranae* is among the most common parasites in *A. mellifera*, with a worldwide distribution (Laurianne et al., 2017). *N. ceranae* infection causes lifespan reduction, energy stress, immunosuppression, apoptosis inhibition, and imbalance of midgut epithelial cell renewal in western honeybees (Antunez et al., 2009; Mayack et al., 2009; Goblirsch et al., 2013; Kurze et al., 2018; Panek et al., 2018). *N. ceranae* can also jeopardize bee hosts together with other biotic or abiotic factors, severely damaging bee health and beekeeping production (Doublet et al., 2015). *N. ceranae* has also been associated with colony collapse disorder (CCD) (Higes et al., 2009; Currie et al., 2010), which severely influences the reproduction and productivity of the colony (Higes et al., 2013; Simeunovic et al., 2014).

To investigate the responses of bee hosts to infection by *N. ceranae*, researchers performed a lot of biochemical and molecular studies and made a series of important progress (Li et al., 2016; Panek et al., 2018; Paris et al., 2018; Huang et al., 2019). For example, Li et al. (2016) silenced the *naked cuticle* gene of *A. mellifera* worker infected by *N. ceranae* using RNAi, resulting in upregulation of the *peptidoglycan recognition protein* encoding gene (*PGRP-S2*) and several antimicrobial peptide encoding genes such as *Abaecin, Apidaecin*, and *Defensin-1*, which reduced the spore number in the host midgut and prolonged the host lifespan. Huang et al. (2019) fed *N. ceranae*-infected western honeybee workers with small interfering RNA (siRNA) targeting the pathogen *Dicer* gene, and found that *N. ceranae* spores were significantly reduced, while the *mucin-2-like* gene of *A. mellifera* was significantly upregulated during 4-6 days after infection, which indicated that *Dicer* was involved in regulating pathogen reproduction and host metabolism. In recent years, on the basis of next-generation sequencing approaches, some omics studies have been conducted to survey the *N. ceranae-response* of *A. mellifera* (Chaimanee et al., 2012; Aufauvre et al., 2014; Huang et al., 2015; Doublet et al., 2016; Badaoui et al., 2017; Fu et al., 2019). Chaimanee et al. (2012) analyzed the expression profiling of immune-related genes in adult worker of *A. mellifera* at 3, 6, and 12 days post infection (dpi) with *N. ceranae*, and observed that humoral immune-related genes such as *Defensin, Abaecin, Apidaecin*, and *Hymenoptaecin* were significantly downregulated at 3 dpi and 6 dpi, whereas the expression level of *Vitellogenin* was not changed, suggestive of immunosuppression for host caused by *N. ceranae*. Our group previously conducted a systematic investigation of the immune responses of *A. m. ligustica* workers to *N. ceranae* invasion based on transcriptome sequencing and bioinformatics, and revealed host cellular and humoral immune responses by analyzing the differential gene expression profiling of *A. m. ligustica* workers challenged by *N. ceranae* (Fu et al., 2019).

As a new member of the noncoding RNA world, circular RNAs (circRNAs) were initially thought to be byproducts of gene transcription without biological function (Cocquerelle et al., 1993). CircRNA has the characteristics of species conservation, stability, and spatiotemporal expression specificity (He et al., 2017). Thanks to the rapid development and application of high-throughput sequencing technology and corresponding bioinformatic algorithms, a batch of circRNAs had been identified in many animal, plant, and microorganism species including *Homo sapiens* (Memczak et al., 2013; Salzman et al., 2013), *Mus musculus* (Memczak et al., 2013), *Triticum aestivum* L (Darbani et al., 2016), *Oryza sativa* (Lu et al., 2015), *Drosophila melanogaster* (Westholm et al., 2015), *Bombyx mori* (Gan et al., 2017), *A. mellifera* (Chen et al., 2019a), *Apis cerana* (Chen et al., 2019b), *N. ceranae* (Guo et al., 2018a), and *Ascosphaera apis* (Guo et al., 2018b). Accumulating experimental and computational evidence indicated the participation of circRNAs in a number of biological processes (He et al., 2017) by regulating the transcription of source genes (Zhang et al., 2013; Li et al., 2015), acting as a “sponge” for microRNA (miRNA) (Xiang et al., 2019), or interacting with various proteins (Ashwal-Fluss et al., 2014). As a result of the covalently closed structure, circRNAs can resist the digestion of RNase R, and hence are ideal biomarkers and therapeutic targets for disease diagnosis and treatment (Meng et al., 2017). More recently, several studies suggested that circRNAs containing internal ribosome entry sites (IRES) or N6-methyladenosine (m^6^A) methylation sites can synthesize biologically active peptides or proteins (Pamudurti et al., 2017; Yang et al., 2017)

Currently, circRNA studies are mainly focused on humans and few model species. For the majority of species, correlational studies on circRNA are lacking. In honeybees, only three circRNA-associated studies have been previously reported (Chen et al., 2019a; Chen et al., 2019b; Thölken et al., 2019). Chen et al. (2019a) predicted 12 211 circRNAs in the ovary of *A. m. ligustica* queen using high-throughput sequencing data, and speculated that differentially expressed circRNA (DEcircRNA) can play an important role in activating queen ovary tissue and oviposition by interacting with miRNAs by analyzing ovaries from virgin queen, egg-laying queen, egg-laying inhibited queen, and egg-laying recovery queen. In our previous work, we conducted deep sequencing of the midguts of *A. m. ligustica* workers at 7 dpi and 10 dpi with *N. ceranae* and corresponding normal midguts using strand-specific cDNA library-based RNA-seq, and identified 10 833 novel circRNAs based on transcriptome data from control groups; it was found that these circRNAs were 15-1 000 nt in length, and the main type of circularization was annotated exon circRNA (Xiong et al., 2018). In humans, chickens, and silkworms, it has been documented that circRNAs are likely to regulate the host response to pathogenic microorganisms (Hu et al., 2018a; Hu et al., 2018b; Shi et al., 2018; Zhang et al., 2019). Hu et al. conducted RNA-seq of normal silkworm midguts and midguts infected by *B. mori* cytoplasmic polyhedrosis virus (BmCPV), screened 294 circRNAs with upregulation and 106 circRNAs with downregulation, and suggested the involvement of corresponding DEcircRNAs in the host response to BmCPV infection by regulating the transcription of sources genes and interacting with target miRNAs (Hu et al., 2018a). However, studies on circRNAs engaged in honeybee parasites/pathogens are lacking until now, and the role that circRNAs play in the western honeybee response to *N. ceranae* invasion remains completely unknown.

To explore the mechanism underlying the immune response of western honeybees to *N. ceranae* infection, we previously performed whole transcriptome sequencing of *N. ceranae*-infected midguts of *A. m. ligustica* workers at 7 dpi and 10 dpi and corresponding uninfected midguts using a combination of strand-specific cDNA library-based RNA-seq and small RNA-seq (sRNA-seq), identified 153 known and 831 novel miRNAs in the midgut of *A. m. ligustica* workers, and analyzed the differential expression pattern of host miRNAs and corresponding target mRNAs during fungal infection, uncovering the mechanism of host immune response mediated by miRNA (Chen et al., 2020); Chen et al. carried out genome-wide identification and characterization of long noncoding RNAs (lncRNAs) in *A. m. ligustica* worker’s midgut, followed by investigation of differentially expressed lncRNAs (DElncRNAs) of host and their regulatory manners such as *cis*-acting effect, *trans*-acting effect, miRNA precursor, and ceRNA network, offering novel insights into host-parasite interaction (Chen et al., 2019c). In this current work, based on our previously obtained high-quality dataset, circRNAs were identified in *N. ceranae*-infected midguts of *A. m. ligustica* workers using bioinformatics followed by analysis of sequence conservation; following function and pathway annotation of source genes of these circRNAs, circRNA-miRNA and circRNA-miRNA-mRNA networks were constructed and analyzed; ceRNA network and circRNAs associated with host cellular and humoral immune were further summarized. To our knowledge, this is the first documentation of circRNA-mediated immune response of honeybee to *N. ceranae* infection, offering a key foundation for clarifying the underlying molecular mechanism of *A. m. ligustica* worker’s *N. ceranae*-response and a new insight into understanding western honeybee-*N. ceranae* interaction.

## 2. Materials and Methods

### 2.1. Biological samples and source of whole transcriptome data

*A. m. ligustica* workers were taken from healthy colonies reared in teaching apiary (119.2369°E, 26.08279°N) of College of Animal Sciences (College of Bee Science), Fujian Agriculture and Forestry University. Clean spores of *N. ceranae* were prepared and kept by honeybee protection laboratory of College of Animal Sciences (College of Bee Science), Fujian Agriculture and Forestry University (Guo et al., 2018a; Guo et al., 2018c; Chen et al., 2019c).

We previously prepared *N. ceranae*-infected midguts of *A. m. ligustica* workers at 7 dpi and 10 dpi and corresponding uninfected midguts (Guo et al., 2018d; Fu et al., 2019). Briefly, (1) newly emerged workers were put into sterilized plastic cages (20 per group) with a feeder containing 50% (w/v) sucrose solution inserted into each box, and incubated at 34 ± 0.5°C for 24 h; (2) after 2 h of starvation treatment, single worker was fixed for artificial inoculation, workers in treatment groups were each fed with 5 μL 50% (w/v) sucrose solution containing 1×10^6^ *N. ceranae* spores (Chen et al., 2019c), while workers in control groups were each fed with 5 μL 50% (w/v) sucrose solution without spores; (3) 24 h after inoculation, workers in both groups were fed with a feeder containing 1 mL 50% (w/v) sucrose solution without spores for 24 h, and the feeder was replaced with a new one every 24 h; each cage was checked daily, and any dead bees were removed; (4) midgut samples of workers (n = 9) in treatment groups and control groups at 7 dpi and 10 dpi were harvested, quickly frozen in liquid nitrogen, and stored at −80°C until deep sequencing. Each treatment group and control group had three biological replicate cages. Midgut samples collected at 7 dpi in control group and treatment group were respectively termed as AmCK1 (AmCK1-1, AmCK1-2, and AmCK1-3) and AmT1 (AmT1-1, AmT1-2, and AmT1-3), while midgut samples collected at 10 dpi were respectively termed as AmCK2 (AmCK2-1, AmCK2-2, and AmCK2-3) and AmT2 (AmT2-1, AmT2-2, and AmT2-3).

In previous study, 12 midgut samples mentioned above were sequenced using strand-specific cDNA library-based RNA-seq technology (Chen et al., 2019c; Chen et al., 2019d). RNA isolation and cDNA library construction were performned according to our previously described method (Chen et al., 2019b; Chen et al., 2019c; Chen et al., 2019d). In brief, (1) total RNA of midgut samples in treatment groups and control groups were respectively extracted using AxyPrep™ Multisource Total RNA Miniprep Kit (TaKaRa, Japan); (2) fragmentation was conducted using divalent cations under elevated temperature in NEBNext First-Strand Synthesis Reaction Buffer (×5); (3) the first-strand cDNA was synthesized using random hexamer primers and reverse transcription, followed by synthesis of second-strand cDNA with DNA polymerase I and RNase H; (4) any remaining overhangs were converted into blunt ends via exonuclease/polymerase activities; (5) the NEBNext adaptor with hairpin loop structures was ligated to prepare for hybridization after adenylation of the 3’ ends of the DNA fragments; (6) the library fragments were purified with AMPure XP Beads (Beckman Coulter, Shanghai, China) to select the insert fragments of 150-200 bp in length, and 3 μL of Uracil-Specific Excision Reagent (USER) Enzyme (NEB, Beijing, China) was then used with the size-selected and adaptor-ligated cDNA at 37°C for 15 min; (7) following PCR amplification with Phusion High-Fidelity DNA Polymerase, universal PCR primers, and an index (X) primer, PCR products were purified (AMPure XP system, Shanghai, China), and the library quality was assessed using an Agilent Bioanalyzer 2100 and real-time PCR; (8) the index-coded samples was clustered on an acBot Cluster Generation System using TruSeq PE Cluster Kitv3-cBot-HS (Illumina, Guangzhou, China) following the manufacturer’s instructions; (9) the constructed 12 cDNA libraries were subjected to deep sequencing on Illumina Hiseq™ 4000 platform by Gene Denovo Biotechnology Co. (Guangzhou, China). Raw reads had been uploaded to the Short Read Archive (SAR) database in NCBI under BioProject number: PRJNA406998. In total, about 273.27 Gb raw reads were generated from RNA-seq, and 271.40 Gb clean reads were obtained after strict quality control; on average, the ratio of clean reads among raw reads in every group was 99.32%, and *Q*20 and *Q*30 were 97.34% and 93.82%, respectively. Hence, the high-quality transcriptome data can be used for identification and investigation of circRNAs, target prediction and analysis, and construction and analysis of regulatory networks in this study.

In another previous work, we sequenced the aforementioned 12 midgut samples using sRNA-seq technology (Chen et al., 2020). RNA extraction and cDNA library construction were performed following our previously described protocol (Chen et al., 2020). Shortly, (1) total RNA of 12 samples was extracted by TRIzol Reagent (Invitrogen, Carlsbad, CA, USA), and DNA contaminants were removed with RNase-free DNase I (TaKaRa, Beijing, China); the purified RNA quantity and quality were checked using Nanodrop 2000 spectrophotometer (Thermo Fisher, Waltham, MA, USA), and the integrity of the RNA samples was evaluated using Agilent 2100 bioanalyzer (Agilent Technologies, Santa Clara, CA, USA) and only values of 28S/18S ≥ 0.7 and RIN ≥ 7.0 were considered qualified for the subsequent small RNA library construction; (2) RNA molecules with a size distribution among 18-30 nt were enriched by agarose gel electrophoresis (AGE) and then ligated with 3’ and 5’ RNA adaptors, followed by reverse transcription and enrichment of fragments with adaptors on both ends were enriched via PCR; (4) the subsequent cDNAs were purified and enriched by 3.5% AGE to isolate the expected size (140-160 nt) fractions and eliminate unincorporated primers, primer dimer products, and dimerized adaptors; (5) the 12 cDNA libraries were sequenced on Illumina sequencing platform (HiSeq™ 2000) using the single-end technology by Gene Denovo Biotechnology Co. (Guangzhou, China). Raw reads derived from sequencing had been submitted to the SAR database in NCBI under BioProject number: PRJNA408312. A total of approximately 23.18 Gb raw reads were produced from sRNA-seq, and ~17.85 Gb clean tags were gained after data quality control; on average, the ratio of clean tags among raw reads was 77.04%, and *Q*20 and *Q*30 were 97.34% and 93.82%, respectively. Therefore, the sRNA data with high quality could be used for prediction of circRNAs target miRNAs and miRNAs target mRNAs, and construction and analysis of circRNA-miRNA and circRNA-miRNA-mRNA networks.

### 2.2. Identification and conservative analysis of circRNAs

CircRNAs were identified following our previously described method (Chen et al., 2019b). Firstly, the clean reads were mapped to the *A. mellifera* genome (assembly Amel_4.5) (The Honeybee Genome Sequencing Consortium, 2006) with TopHat software (Kim et al., 2013). Next, 20 nt from the 5’ and 3’ ends of unmapped reads were extracted and aligned independently to reference sequences using Bowtie2 (Langmead et al., 2009). Ultimately, the unmapped anchor reads were submitted to find_circ (Memczak et al. 2013) for the identification of circRNA following criteria: breakpoints = 1; edit ≤ 2; n uniq > 2; anchor overlap ≤ 2; best qual A > 35 or best qual B > 35; n uniq > int (samples/2); circRNA length < 100 kb.

To investigate the conservation of circRNAs of *A. m. ligustica* and other species, sequences of source genes of circRNAs identified in this study were aligned to those previously identified in *Apis cerana cerana* (Chen et al., 2019b) and human (*H. sapiens*) (Ashwal-Fluss et al., 2014; Rybak-Wolf et al., 2015).

### 2.3. Analysis of DEcircRNAs and their source genes

For each circRNA, the expression level was calculated using RPM = 10^6^C/N (C is the number of back-spliced junction reads that uniquely aligned to a circRNA, and N is the total number of back-spliced junction reads). DESeq software (Wang et al., 2010) was used to identify significant DEcircRNAs in AmCK1 vs AmT1 and AmCK2 vs AmT2 comparison groups following the threshold of |log2(Fold change)| ≥ 1 and *P* ≤ 0.05.

CircRNA can regulate source gene expression via interaction with RNA polymerase II, U1 micronucleoprotein or gene promoter (Zhang et al., 2013; Li et al., 2015). In the present study, source genes of DEcircRNAs were predicted by alignment of anchors reads at both ends to *A. mellifera* genome (assembly Amel_4.5) (The Honeybee Genome Sequencing Consortium, 2006) using Bowtie tool (Langmead et al., 2009). Gene Ontology (GO) term analysis for source genes was performed with DAVID gene annotation tool (http://david.abcc.ncifcrf.gov/) (Huang et al., 2007). Two-sided Fisher’s exact test was carried out to classify the GO category, while the FDR was calculated to correct the *P* value (Jung, 2014). GO terms with a *P* value <0.05 were considered to be statistically significant. In addition, pathway analysis was performed through annotating source genes to Kyoto Encyclopedia of Genes and Genomes (KEGG) database (http://www.genome.jp/kegg/) (Du et al., 2014). The significance threshold was defined by a *Q* value < 0.05.

### 2.4. Target prediction and regulatory network construction of DEcircRNAs

Target miRNAs of DEcircRNAs were predicted using TargetFinder software (Allen et al., 2009). Following the cutoff of *P* ≤ 0.05 and free energy ≤ 35, potential target miRNAs were further extracted to construct DEcircRNA-miRNA regulatory network. Target mRNAs of miRNAs targeted by DEcircRNAs were further predicted with TargetFinder software, followed by construction of DEcircRNA-miRNA-mRNA regulatory network. The regulatory networks were visualized by Cytoscape software (Smoot et al., 2011). Moreover, target mRNAs within DEcircRNA-miRNA-mRNA network were annotated to GO and KEGG databases utilizing Blast tool. Based on function and pathway annotations and our previous results (Fu et al., 2019; Chen et al., 2019c; Chen et al., 2020), DEcircRNA-differentially expressed miRNA (DEmiRNA)-differentially expressed mRNA (DEmRNA) regulatory network relative to host cellular and humoral immune was constructed and then visualized with Cytoscape.

### 2.5. PCR of novel circRNAs

Three circRNAs (novel_circ_004065, novel_circ_002199, and novel_circ_005784) were randomly selected from specific circRNAs in AmT1 and AmT2 for molecular verification. Theoretically, for circRNA, convergent primers should amplify products from both the cDNA template and the gDNA template, but divergent primers should only amplify products from the cDNA template (Chen et al., 2019b). According to the method described by Chen et al. (2019b) and Xiong et al. (2018), specific convergent primers and divergent primers (shown in Table S1) for above-mentioned circRNAs were designed with DNAMAN 8 software (Lynnon Biosoft, USA), and synthesized by Shanghai Sangon biological co., Ltd. The total RNA of AmT1 and AmT2 were respectively extracted with AxyPre RNA extraction kit (Axygen, China) and divided into two portions. One portion was digested with 3 U/mg RNase R (Geneseed, China) at 37°C for 15 min to remove linear RNA, and then synthesize the first-strand cDNA by transcription with random primers, and the other was subjected to reverse transcription using Oligo (dT)_18_ as a template to synthesize the first-strand cDNA. Genomic DNA (gDNA) of AmT1 and AmT2 were respectively isolated using the AxyPre DNA extraction kit (Axygen, China). The equimolar mixture of AmT1 and AmT2 cDNA was used as templates for PCR, which was conducted on a T100 thermocycler (Bio-Rad, USA) in a 20 μL reaction volume containing 1 μL of template, 10 μL of Mixture (TaKaRa, Japan), 1 μL upstream primers (10 μmol/L) and 1 μL downstream primers (10 μmol/L), and 7 μL of dH_2_O. PCR conditions were as follows: pre-denaturation at 94°C for 5 min, followed by 36 cycles of denaturation at 94°C for 50 s, an appropriate annealing temperature (according to the melting temperature of the primer) for 30 s, and extension at 72°C for 1 min, and final extension at 72°C for 5 min. The PCR products were detected on 1.5% agarose gel electrophoresis and sequenced by Sanger sequence (Sangon biological co., Ltd, Shanghai).

### 2.6. Stem-loop RT-PCR verification of miRNA

Ten miRNAs within DEcircRNA-miRNA-mRNA regulatory network were randomly selected for molecular validation, including mir-30-x, mir-451-x, mir-29-y, ame-miR-3720, mir-21-x, mir-146-x, mir-143-y, mir-101-y, mir-462-x and mir-7975-y. Specific Stem-loop primers, specific upstream primers, and universal downstream primers (shown in Table S2) of aforementioned miRNAs were designed using DNAMAN 8 software (Lynnon Biosoft, USA). Total RNA of AmCK1, AmT1, AmCK2 and AmT2 were respectively extracted using Eastep^®^ Super Total RNA extraction kit (Promega, China), followed by synthesis of corresponding cDNA with Stem-loop primers. PCR reaction was carried out in a 20 μL reaction volume containing 1 μL cDNA template, 1 μL upstream primers (10 μmol/L), 1 μL downstream primers (10 μmol/L), 7 μL dH_2_O, and 10 μL PCR Mixture (TaKaRa, Japan). PCR reaction was conducted on a T100 thermocycler (Bio-Rad, Hercules, CA, USA) following the parameters: pre-denaturation at 94°C for 5 min; 36 cycles of denaturation at 94°C for 50 s, annealing at 56°C for 30 s, and elongation at 72°C for 1 min; final elongation at 72°C for 10 min. PCR products were examined on 2% agarose gel electrophoresis.

### 2.7. Real-time quantitative PCR (RT-qPCR) validation of DEcircRNAs

Eight DEcircRNAs were randomly selected for RT-qPCR verification, including novel_circ_000705, novel_circ_001195, novel_circ_011173, novel_circ_006925, novel_circ_012352 and novel_circ_012316 from AmCK1 vs AmT1, and novel_circ_007686 and novel_circ_011500 from AmCK2 vs AmT2. Specific divergent primers and convergent primers (Table S1) for DEcircRNAs were designed with DNAMAN 8 software and molecular verification according to the above-mentioned method. Total RNA of AmCK1, AmT2, AmCK2, and AmT2 was respectively extracted and divided into two parts. One part was digested with 3 U/mg RNase R (Geneseed, China) at 37°C for 15 min followed by reverse transcription with random primer, and the resulting cDNA was used as template for DEcircRNAs; the other was reversely transcribed with Oligo (dT)_18_, and the resulting cDNA was used as template for reference gene (*actin*). The reaction system was 20 μL, containing 1 μL forward primer (10 μmol·L-1), 1 μL reverse primer (10 μmol·L-1), 1 μL cDNA template, 10 μL SYBR Green Dye, and 7 μL DEPC water. The reaction was conducted on a QuantStudio3 real-time PCR instrument (ABI, USA), and each reaction was performed in triplicate following the instruction of SYBR Green Dye Kit (Vazyme, China). The reaction conditions were set as: pre-denaturation at 94°C for 5 min, denaturation at 94°C for 50 s, extension at 60°C for 30 s, a total of 45 cycles. The relative expression of DEcircRNAs was calculated using the 2^-ΔΔCt^ method (Livak and Schmittgen, 2001) and presented as relative expression levels from three biological replicates and three parallel replicates, followed by visualization with GraphPad Prism 7 software (GraphPad, USA).

## 3. Results

### 3.1. Processing and quality control of the dataset

On average, there were 236 067 761 and 326 923 515 anchors reads in AmT1 and AmT2, respectively, and more than 16 349 641 and 18 445 112 anchors reads could be mapped to the genome of *A. mellifera* (Table 1). Additionally, Pearson correlations between the different biological replicas within both AmT1 and AmT2 groups were above 0.968 (Fig. S1). Our previous results showed that there were an average of 180 533 501 and 182 634 149 anchors reads in the AmCK1 and AmCK2 groups, respectively; 18 794 099 and 18 687 848 could be mapped to the reference genome (Xiong et al., 2018); Pearson correlation coefficients between the different biological replicates were above 0.950 (Guo et al., 2018d). The results suggested that the sequencing data and sample preparation in this work were reasonable and reliable.

**Table 1.**
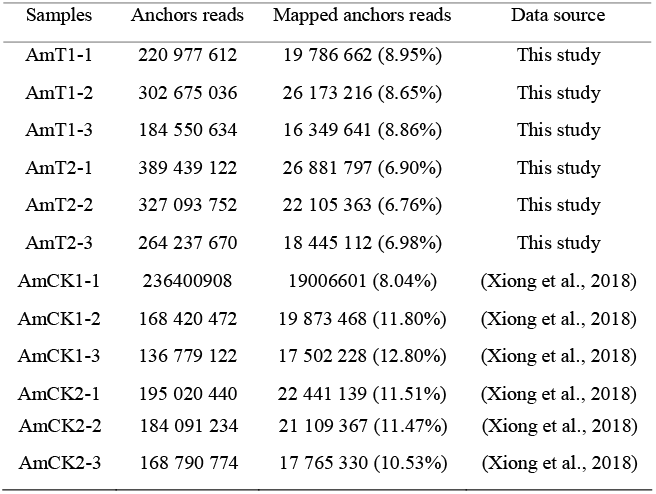
Mapping of anchors reads to the reference genome of *A. mellifera*

### 3.2 Quantity, expression, conservation of A. m. ligustica circRNAs

Here, 8 199 and 8 711 circRNAs were identified in the AmT1 and AmT2 groups, respectively. In our previous work, 6 530 and 6 289 circRNAs were identified in AmCK1 and AmCK2 groups, respectively (Xiong et al., 2018). In total, 14 909 *A. m. ligustica* circRNAs were identified using a combination of datasets from *N. ceranae*-infected and uninfected groups; 1 398, 1 696, 1 019, and 1 871 circRNAs were specifically transcribed in each group (Fig. 1A). Moreover, distinct PCR products with the expected size were amplified with the divergent primers (Fig. 1B and Fig. 1C). This confirmed that *A. m. ligustica* circRNAs identified in this study were credible.

**Fig. 1.**
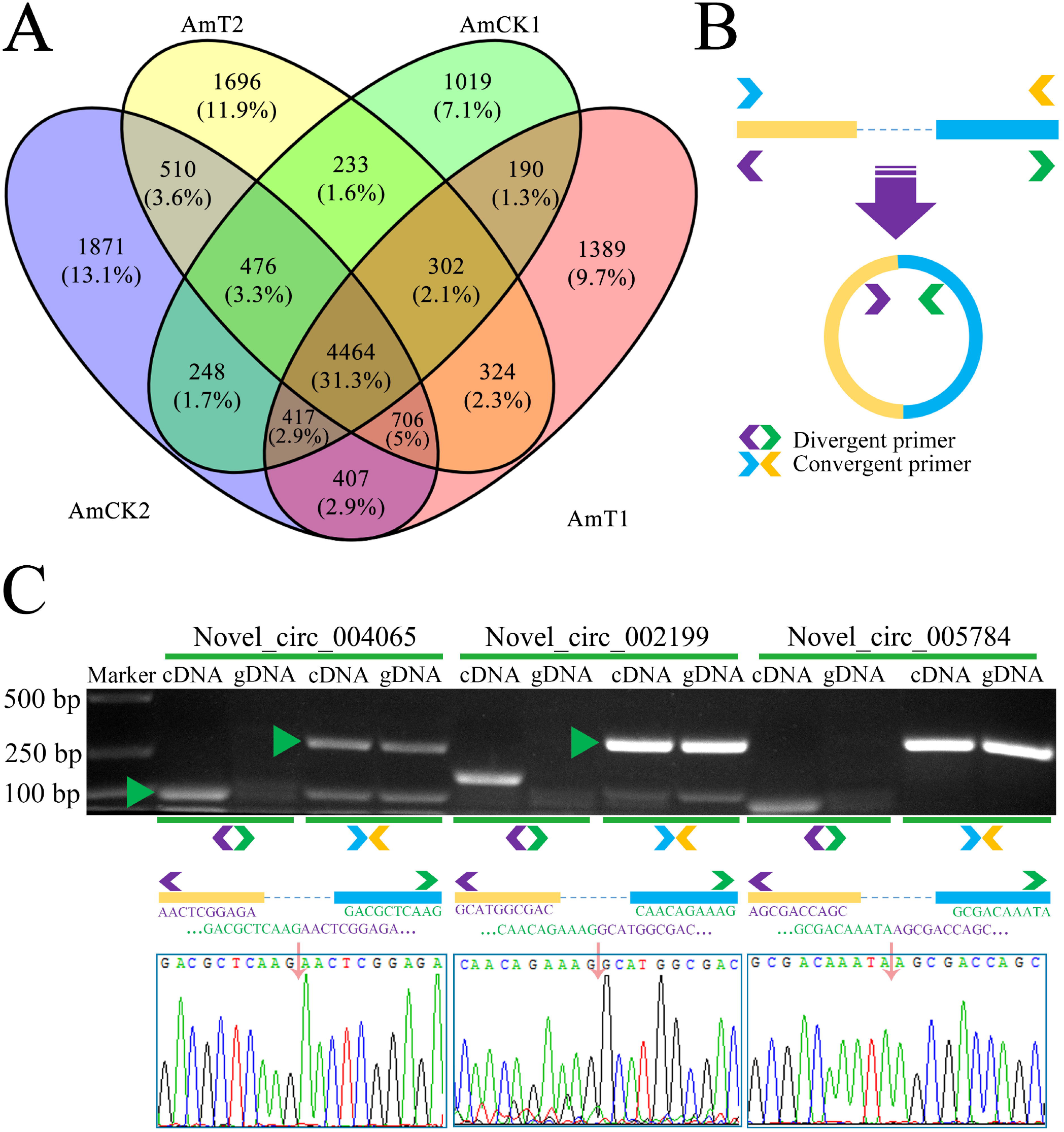
Venn analysis and molecular validation of *A. m. ligustica* circRNAs. Venn analysis of circRNAs identified in four groups (A). A schematic diagram showing the designing of convergent primers and divergent primers for circRNAs (B). Agarose gel electrophoresis and Sanger sequence for amplified products from novel_circ_004065, novel_circ_002199, and novel_circ_005784 with divergent primers (C).

The source gene sequences of 14 909 identified *A. m. ligustica* circRNAs were aligned with those of *A. c. cerana* (Chen et al., 2019b) and *H. sapiens* (Ashwal-Fluss et al., 2014; Rybak-Wolf et al., 2015) circRNAs, and the results demonstrated that only 20 (0.13%) *A. m. ligustica* circRNAs had homology with *H. sapiens* circRNAs, while as many as 10 226 (68.59%) *A. m. ligustica* circRNAs were homologous to *A. c. cerana* circRNAs (Fig. 2). Additionally, there were 16 (0.11%) conservative circRNAs in these three species (Fig. 2).

**Fig. 2.**
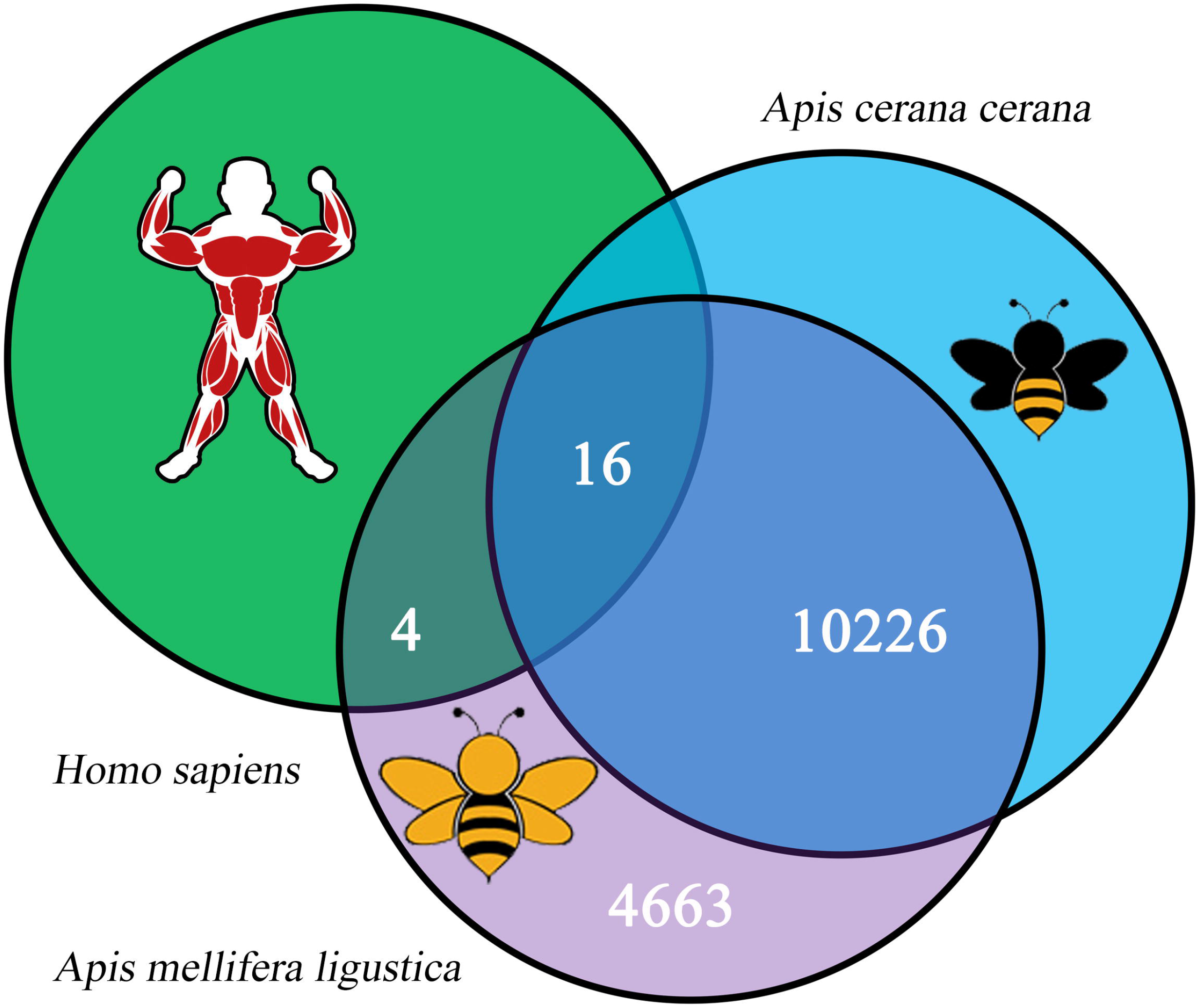
Sequence conservation of 14909 *A. m. ligustica* circRNAs.

### 3.3. DEcircRNAs in the midguts of A. m. ligustica workers response to N. ceranae infection

The overall expression level of circRNAs in AmCK1 was slightly higher than that of AmT1, while circRNAs in AmCK2 had a slightly lower overall expression level than those in AmT2 (Fig. 3A). In total, 168 DEcircRNAs were screened out in the AmCK1 vs AmT1 comparison group, including 61 upregulated circRNAs and 107 downregulated circRNAs (Fig. 3B, see also Table S3). Among these DEcircRNAs, the top three circRNAs with the highest upregulation were novel_circ_004821 [log_2_(Fold change) = 16.1509], novel_circ_002611 [log2(Fold change) = 16.0817], and novel_circ_000878 [log_2_(Fold change) = 16.0385]; the most downregulated circRNA was novel_circ_014612 [log_2_(Fold change) = −16.8472], followed by novel_circ_012545 [log_2_(Fold change) = −16.8103] and novel_circ_005159 [log_2_(Fold change) = −16.7233] (Table S3). In the AmCK2 vs AmT2 comparison group, 306 DEcircRNAs including 143 upregulated circRNAs and 163 downregulated circRNAs were identified (Fig. 3B, see also Table S4). Among them, novel_circ_002577 [log_2_(Fold change) = 16.2244] was the most up-regulated, followed by novel_circ_011847 [log_2_(Fold change) = 16.0676] and novel_circ_007020 [log_2_(Fold change) = 16.8271]; the most down-regulated circRNAs were novel_circ_006432 [log_2_(Fold change) = −17.3774], novel_circ_011107 [log_2_(Fold change) = −16.1564], and novel_circ_002456 [log_2_(Fold change) = −15.8598] (Table S4). Additionally, Venn analysis indicated that nine upregulated circRNAs and 10 downregulated circRNAs were shared by both comparison groups (Fig. 3C); 54 upregulated circRNAs and 97 downregulated ones were unique for AmCK1 vs AmT1, whereas 134 upregulated circRNAs and 155 downregulated ones were unique for AmCK2 vs AmT2 (Fig. 3C).

**Fig. 3.**
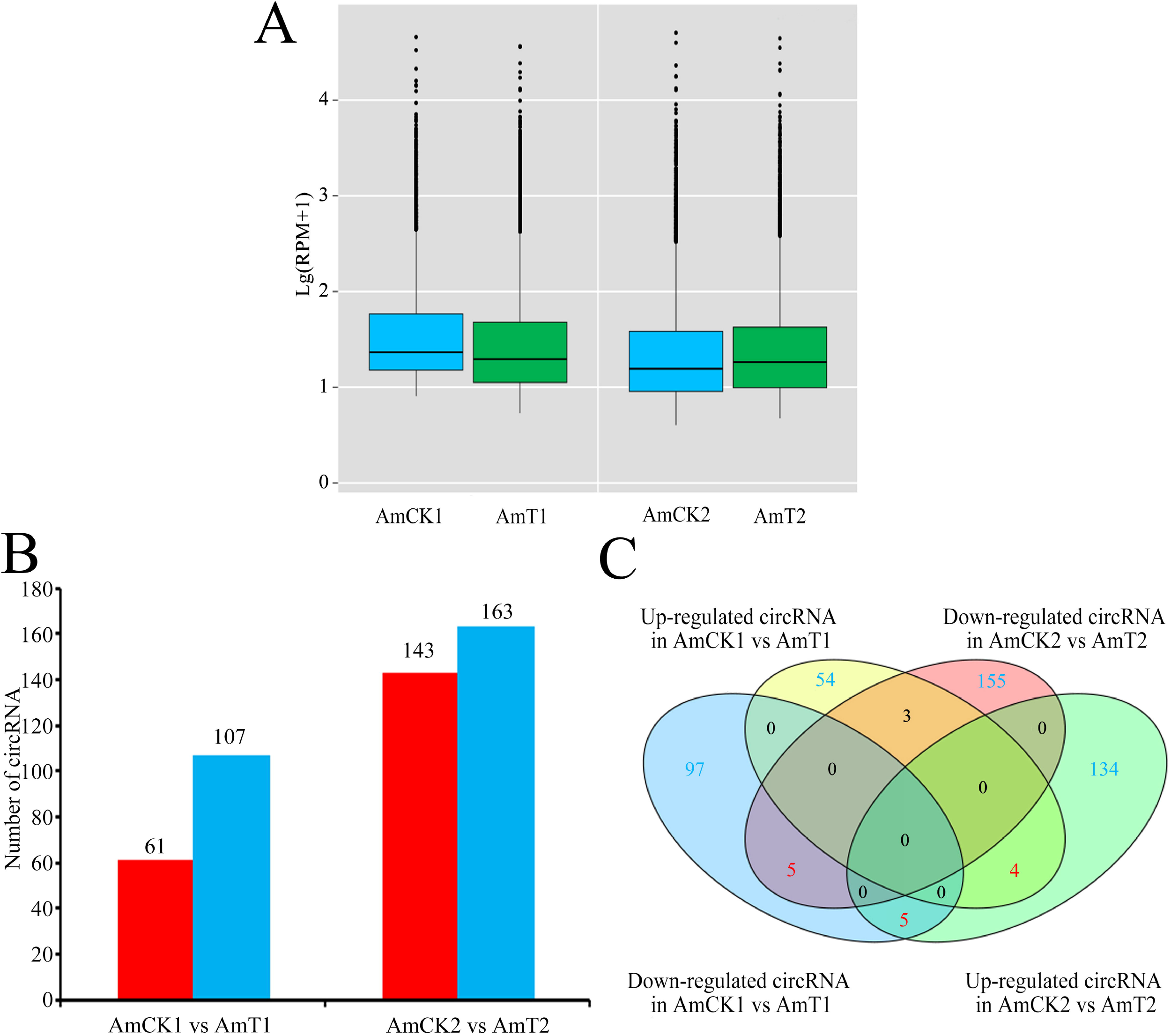
Differential expression profile of circRNAs in *N. ceranae-infected* and uninfected midguts of *A. m. ligustica* workers. Boxplots showing the overall expression levels of circRNAs identified in four groups. Number of DEcircRNAs in two comparison groups; red columns indicate up-regulated circRNAs, while blue columns indicate down-regulated ones (B). Venn analysis of DEcircRNAs (C).

Furthermore, back-splicing sites of eight DEcircRNAs were validated using PCR with divergent primers and Sanger sequencing(Fig. 4A); additionally, RT-qPCR results suggested that the expression trend of eight DEcircRNAs was in accordance with that detected in sequencing data. This confirmed the reliability of the transcriptome dataset used in the present study (Fig. 4B).

**Fig. 4.**
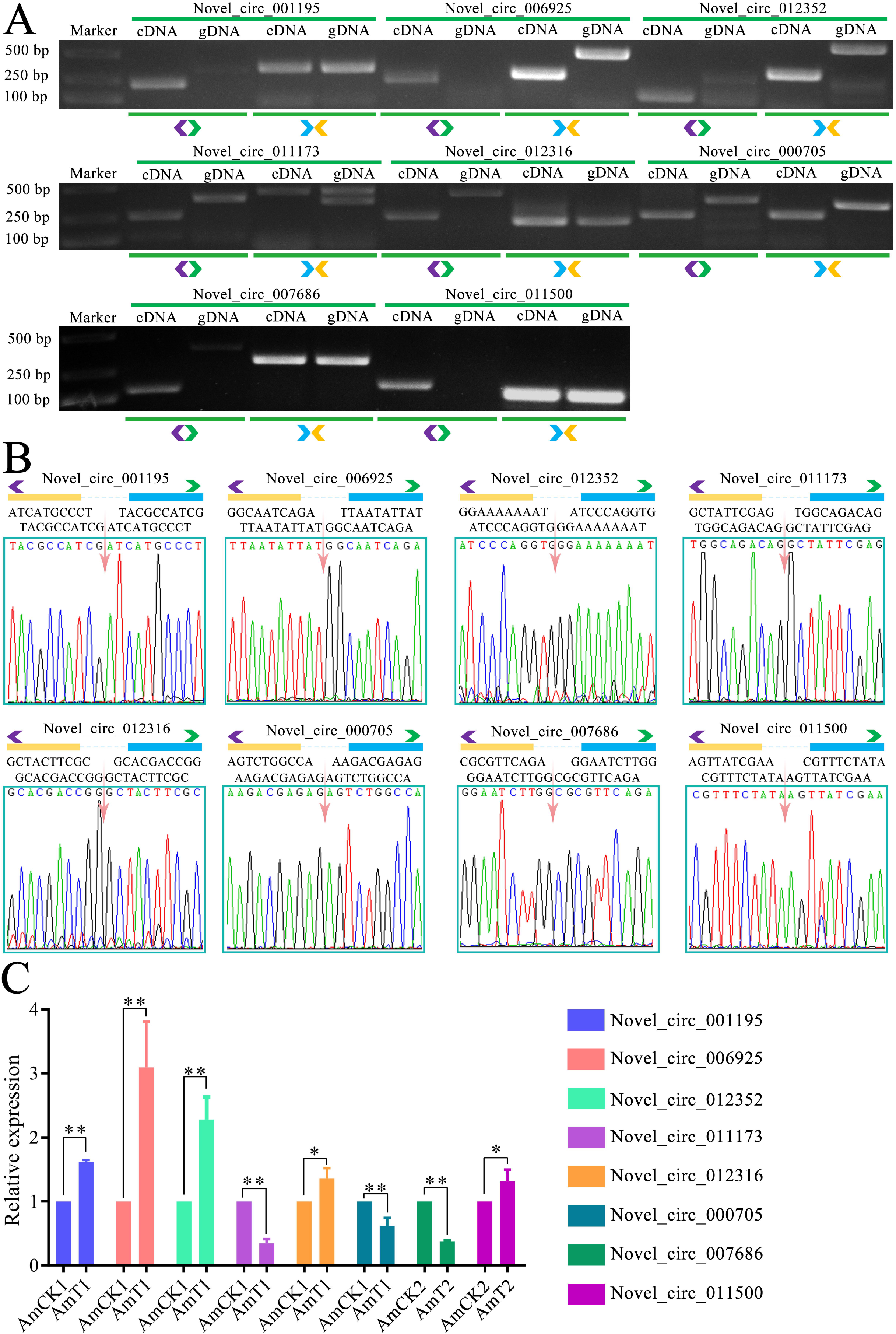
PCR, Sanger sequencing, and RT-qPCR validation of DEcircRNAs. Agarose gel lectrophoresis of products from PCR amplification of eight DEcircRNAs with specific divergent primers (A). Sanger sequencing of amplified fragments from eight DEcircRNAs (B). RT-qPCR results of eight DEcircRNAs, “*” indicates *P* < 0.05, “**” indicates *P* < 0.01 (C).

### 3.4. Function and pathway annotation of source genes of DEcircRNAs

Based on GO database annotation, it’s found that source genes of DEcircRNAs in AmCK1 vs AmT1 comparison group could be annotated to 27 functional terms relative to biological process, cellular component, and molecular function. In biological process category, the largest subcategory was cellular process (21), followed by single-organism process (18), metabolic process (17), localization (eight), and biological regulation (six) (Fig. 5A, see also Table S5); in cellular component category, the most abundant groups were cell part (ten), cell (ten), membrane (eight), membrane part (seven), and organelle (seven) (Fig. 5A, see also Table S5); in molecular function category, to the top five categories were binding (31), catalytic activity (20), transporter activity (four), nucleic acid binding transcription factor activity (two), and molecular transducer activity (one) (Fig. 5A, see also Table S5). In AmCK2 vs AmT2 comparison group, source genes of DEcircRNAs can be annotated to 35 functional terms, including 15 biological process-associated terms such as cellular process (63), single-organism process (55), and metabolic process (53) (Fig. 5B, see also Table S6); 12 cellular component-associated terms such as membrane (27), membrane part (26), cell (26), and cell part (26) (Fig. 5B, see also Table S6); and eight molecular function-associated terms such as binding (77), catalytic activity (47), and nucleotide binding transcription factor activity (11) (Fig. 5B, see also Table S6).

**Fig. 5.**
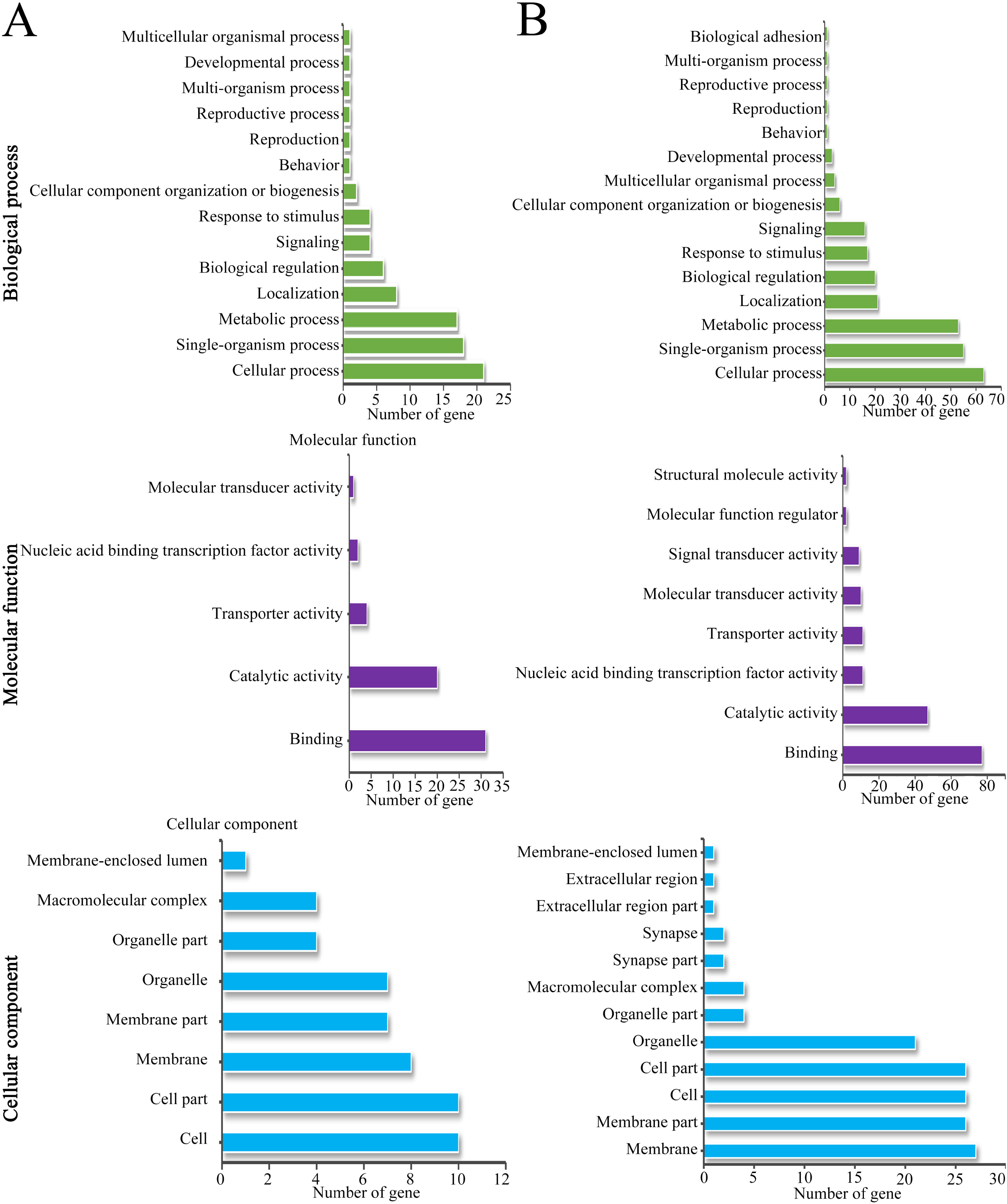
Functional annotation of DEcircRNAs’ source genes in AmCK1 vs AmT1 (A) and AmCK2 vs AmT2 (B) comparison groups.

*N. ceranae* infection can not only result in damage of structure and renewal of honeybee’s gut tissue (Panek et al., 2018), but also influence host immune response (Antunez et al., 2009). Further analysis revealed that 21 (63) and two (six) source genes of DEcircRNAs in AmCK1 vs AmT1 (AmCK2 vs AmT2) comparison group were engaged in cell renewal-related terms including cellular process and cell component organization or biosynthesis (Fig. 5, see also Table S5 and Table S6); additionally, ten (26), ten (26), eight (27), seven (26), seven (21), four (four), one (one) source genes were involved in cell structure-associated terms such as cell part, cell, membrane, membrane part, and membrane-enclosed lumen (Fig. 5, see also Table S5 and Table S6); moreover, four (six) source genes were annotated to response to stimulus, a functional term relative to host immune defense system (Fig. 5, see also Table S5 and Table S6).

### 3.6. KEGG pathway annotation of source genes of DEcircRNAs

KEGG database annotation result demonstrated that source genes of DEcircRNAs in AmCK1 vs AmT1 comparison group were associated with 33 pathways, including endocytosis (four), FoxO signaling pathway (three), phagosome (three), galactose metabolism (two), and arginine and proline metabolism (two) (Fig. 6A, see also Table S7); while those in AmCK2 vs AmT2 comparison group were involved in 43 pathways, such as endocytosis (eight), Hippo signaling pathway (six), starch and sucrose metabolism (four), insulin resistance (four), and purine metabolism (four) (Fig. 6B, see also Table S8).

**Fig. 6.**
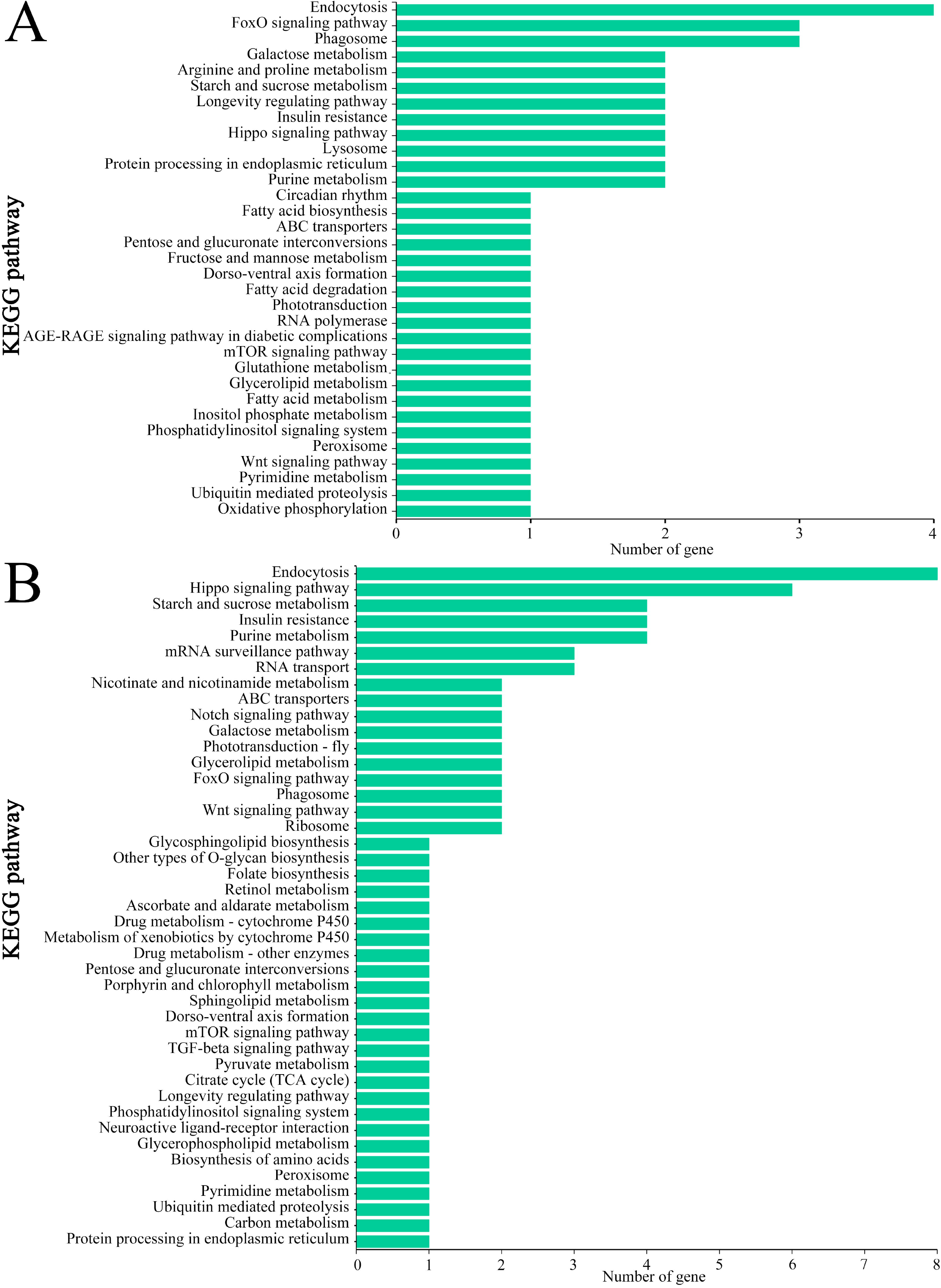
KEGG pathway annotation of source genes of DEcircRNAs in AmCK1 vs AmT1 (A) and AmCK2 vs AmT2 (B) comparison groups.

Further analysis was performed to explore pathways relative to cell renewal, carbohydrate metabolism, and energy metabolism, and the result suggested that in both comparison groups mentioned above, two (one) and six (two) source genes of DEcircRNAs were involved in Hippo signaling pathway and Wnt signaling pathway (Fig. 6, see also Table S7 and Table S8); additionally, two, two, one, and one source genes of DEcircRNAs in AmCK1 vs AmT1 were annotated to galactose metabolism, starch and sucrose metabolism, as well as fructose and mannose metabolism; while only one source gene was annotated to oxidative phosphorylation, a key energy metabolism-related pathway in eukaryotes (Fig. 6A, Table S7). There were two and four source genes of DEcircRNAs in AmCK2 vs AmT2 involved in galactose metabolism as well as starch and sucrose metabolism (Fig. 6B, see also Table S8).

Moreover, pathways relative to host immune were examined, and the result indicated that four, three, and two source genes of DEcircRNAs in AmCK1 vs AmT1 comparison group were annotated to several cellular immune pathways such as endocytosis, phagosome, and lysosome (Table 2); while eight, two, one and one source genes of DEcircRNAs in AmCK2 vs AmT2 comparison group were engaged in endocytosis, phagosome, ubiquitin-mediated proteolysis, and metabolism of xenobiotics by cytochrome P450 (Table 2). Intriguingly, source gene of DEcircRNA in both comparison groups was observed to annotate to only one humoral immune pathway, FoxO signaling pathway (Fig. 6, see also Table 2).

**Table 2.**
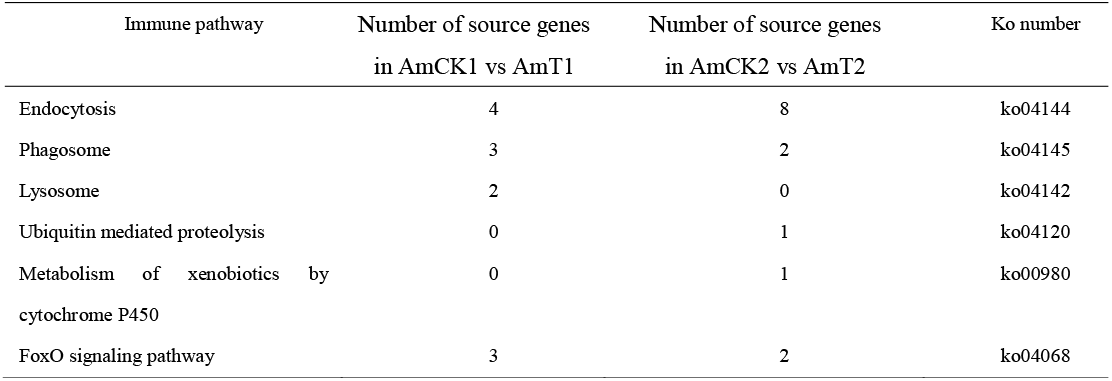
Source genes of DEcircRNAs engaged in host cellular and humoral immune pathways

### 3.7. Analysis of the DEcircRNA-miRNA regulatory network involved in A. m. ligustica workers’ midgut response to N. ceranae

A total of 82 target miRNAs of 122 DEcircRNAs in the AmCK1 vs AmT1 comparison group were predicted, among which ovel_circ_011088, novel_circ_013731, and novel_circ_009951 had the most targets, amounting to 48, 37, and 22, respectively (Fig. 7A, see also Table S9). In addition, mir-8503-x could be targeted by as many as 16 DEcircRNAs, while novel-m0007-5p and mir-151-x could be targeted by 14 and 11 DEcircRNAs, respectively (Fig. 7A, see al Table S9). In the AmCK2 vs AmT2 comparison group, 106 target miRNAs of 234 DEcircRNAs were identified, and novel_circ_011577, novel_circ_002577, and novel_circ_012916 had the most targets, reaching 53, 24, and 24, respectively (Fig. 7B, see also Table S10); additionally, mir-8503-x (24 DEcircRNAs), ame-miR-3747b (22 DEcircRNAs), and ame-miR-981 (21 DEcircRNAs) could be targeted by the most DEcircRNAs (Fig. 7B, see also Table S10). Furthermore, Stem-loop RT-PCR confirmed the expression of ten miRNAs within DEcircRNA-miRNA regulatory networks (Fig. 7C).

**Fig. 7.**
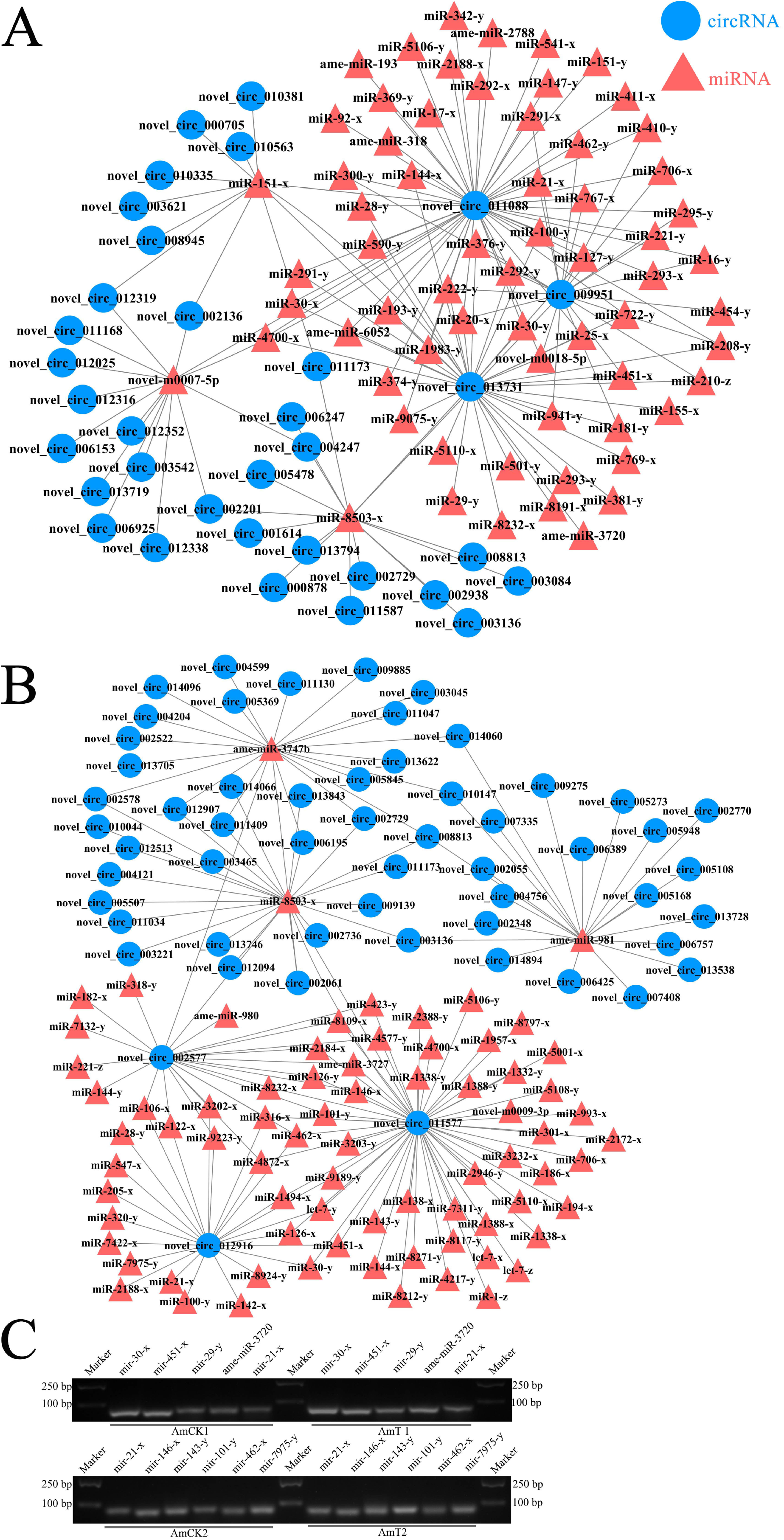
DEcircRNA-miRNA regulatory network. Regulatory networks of DEcircRNAs and their target -miRNAs in AmCK1 vs AmT1 (A). Regulatory networks of DEcircRNAs and their target -miRNAs in AmCK2 vs AmT2 (B). Stem-loop PCR validation of DEcircRNA target miRNAs (C).

### 3.8 Investigation of the ceRNA regulatory network associated with N. ceranae-response of A. m. ligustica workers’ midguts

Target mRNAs of DEcircRNA-targeted miRNAs were predicted followed by construction and investigation of the ceRNA regulatory network. The results demonstrated that 86 DEcircRNAs in AmCK1 vs AmT1 could bind to 75 miRNAs, further targeting 215 mRNAs (Table S11); while 178 DEcircRNAs in AmCK2 vs AmT2 could link to 103 miRNAs, further targeting 305 mRNAs (Table S12).

Functional annotation indicated that mRNAs within ceRNA networks in the midgut of *A. m. ligustica* workers at 7 dpi with *N. ceranae* infection were engaged in 33 functional terms, including two cell renewal-associated terms such as cellular process and cellular component organization or biogenesis, seven cell structure-associated terms such as cell and membrane, as well as two immune-associated terms such as response to stimulus and cell killing (Fig. 8A, see also Table S13); mRNAs within ceRNA regulatory network in the worker’s midgut at 10 dpi were involved in 28 functional terms, including two cell renewal-related functional terms such as cellular process and cellular component organization or biogenesis, six cell structure-related functional terms such as cell part and membrane part, and one immune-related term (response to stimulus) (Fig. 8B, see also Table S14).

**Fig. 8.**
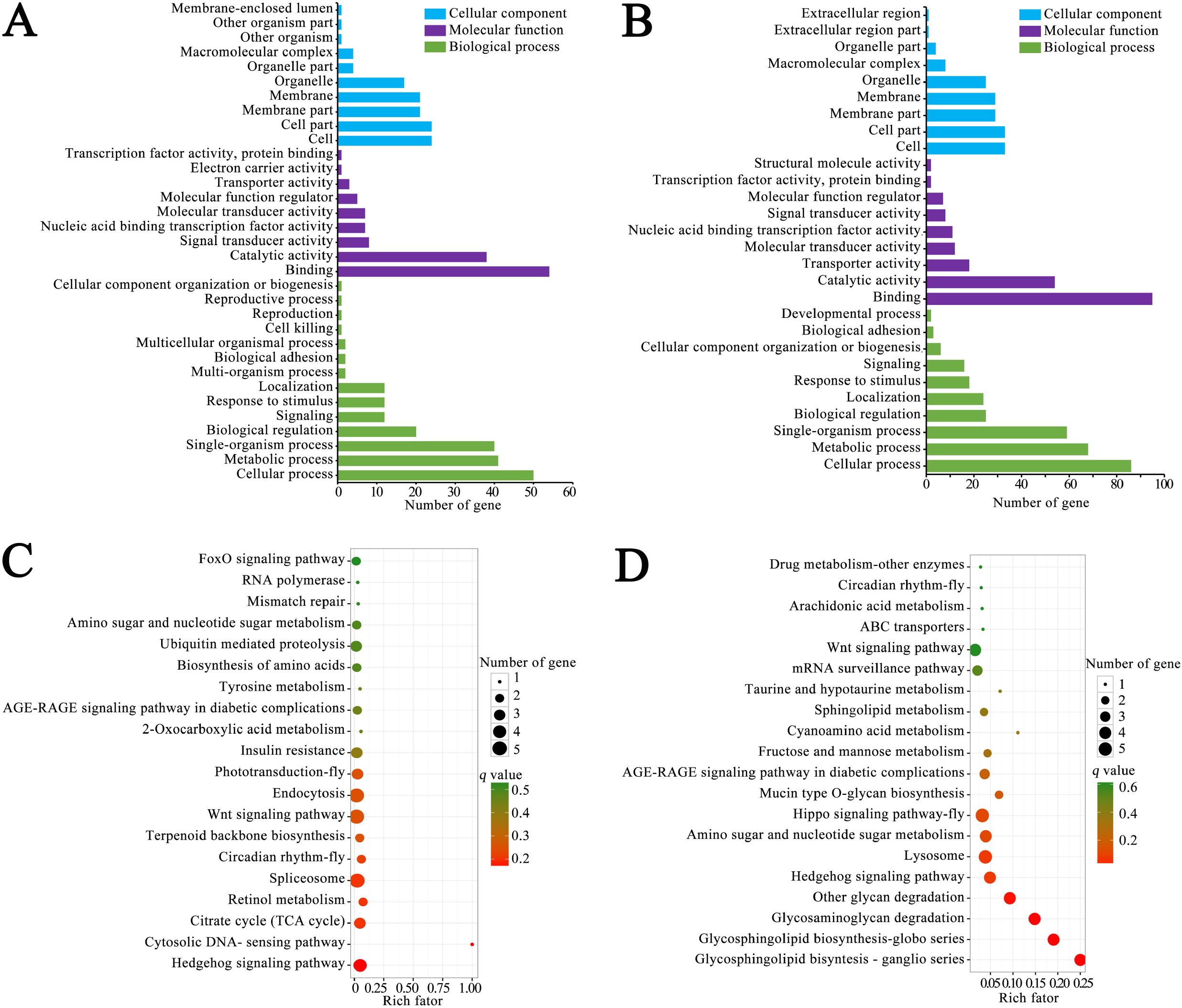
Function and pathway annotations of target mRNAs within ceRNA regulatory networks. A-B: GO database annotation of target mRNAs of DEcircRNA-targeted miRNAs in AmCK1 vs AmT1 and AmCK2 vs AmT2 (A-B). KEGG database annotation of target mRNAs of DEcircRNA-targeted miRNAs in AmCK1 vs AmT1 and AmCK2 vs AmT2 (C-D).

Moreover, mRNAs within the ceRNA regulatory network in AmCK1 vs AmT1 could also be annotated to 41 pathways, including two gut development-related pathways(Hippo signaling pathway and Wnt signaling pathway), two sugar metabolism-related pathways (amino sugar and nucleotide sugar metabolism as well as galactose metabolism), and one energy metabolism-related pathway (oxidative phosphorylation); additionally, three targets were engaged in cellular immune pathways including endocytosis, phagosomes, and ubiquitin-mediated proteolysis, while two were associated with humoral immune pathways such as the FoxO signaling pathway and the MAPK signaling pathway (Fig. 8C, see also Table S15). In the AmCK2 vs AmT2 comparison group, target mRNAs can be annotated to 47 pathways, including Hippo signaling pathway, Wnt signaling pathway, amino sugar and nucleotide sugar metabolism, fructose and mannose metabolism, pentose phosphate pathway, glycolysis, gluconeogenesis, endocytosis, lysosome, ubiquitin-mediated proteolysis, FoxO signaling pathway, and MAPK signaling pathway (Fig. 8D, see also Table S16).

Based on our previous studies on DEmRNAs, DEmiRNAs, and DElncRNAs involved in *A. m. ligustica* worker midgut responses to *N. ceranae* infection (Fu et al., 2019; Chen et al., 2019c; Chen et al., 2020), we further analyzed DEcircRNA-DEmiRNA-DEmRNA regulatory networks associated with host cellular and humoral immune response, and the resultsdemonstrated that 16 DEcircRNAs in AmCK1 vs AmT1 can link to 15 DEmiRNAs, further targeting 10 DEmRNAs relative to endocytosis, ubiquitin-mediated proteolysis, phagosomes, FoxO signaling pathway, and MAPK signaling pathway (Fig. 9B, see also Table 3), whereas three DEcircRNAs in AmCK2 vs AmT2 could bind to 26 DEmiRNAs, further targeting 10 DEmRNAs relative to lysosome, ubiquitin-mediated proteolysis, endocytosis, MAPK signaling pathway, and FoxO signaling pathway (Fig. 9C, see also Table 3).

**Fig. 9.**
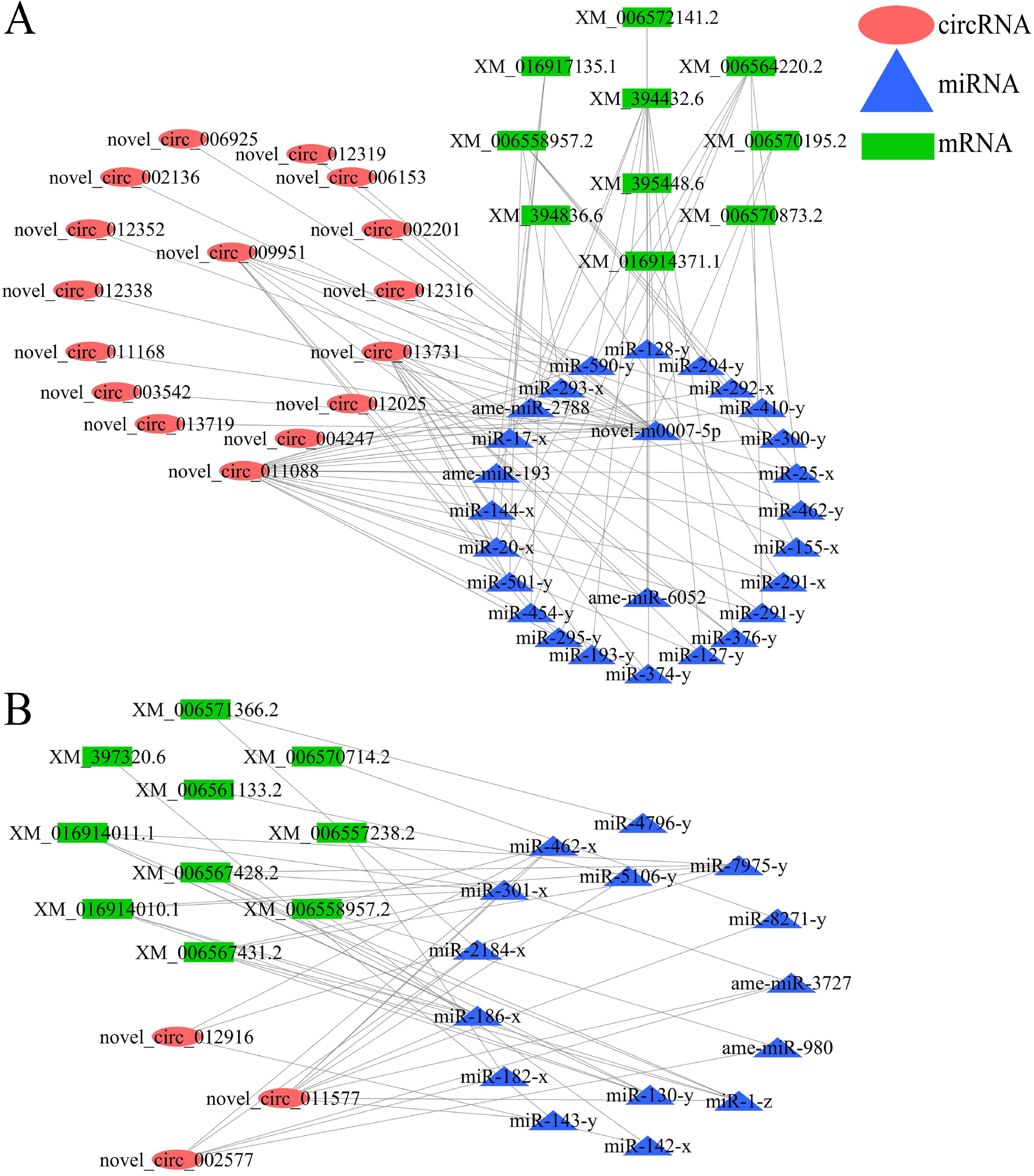
CeRNA regulatory networks relative to host cellular and humoral immune. Cellular and humoral immune-associated ceRNA network of DEcircRNAs in AmCK1 vs AmT1 (A). Cellular and humoral immune-associated ceRNA network of DEcircRNAs in AmCK2 vs AmT2 (B).

**Table 3.**
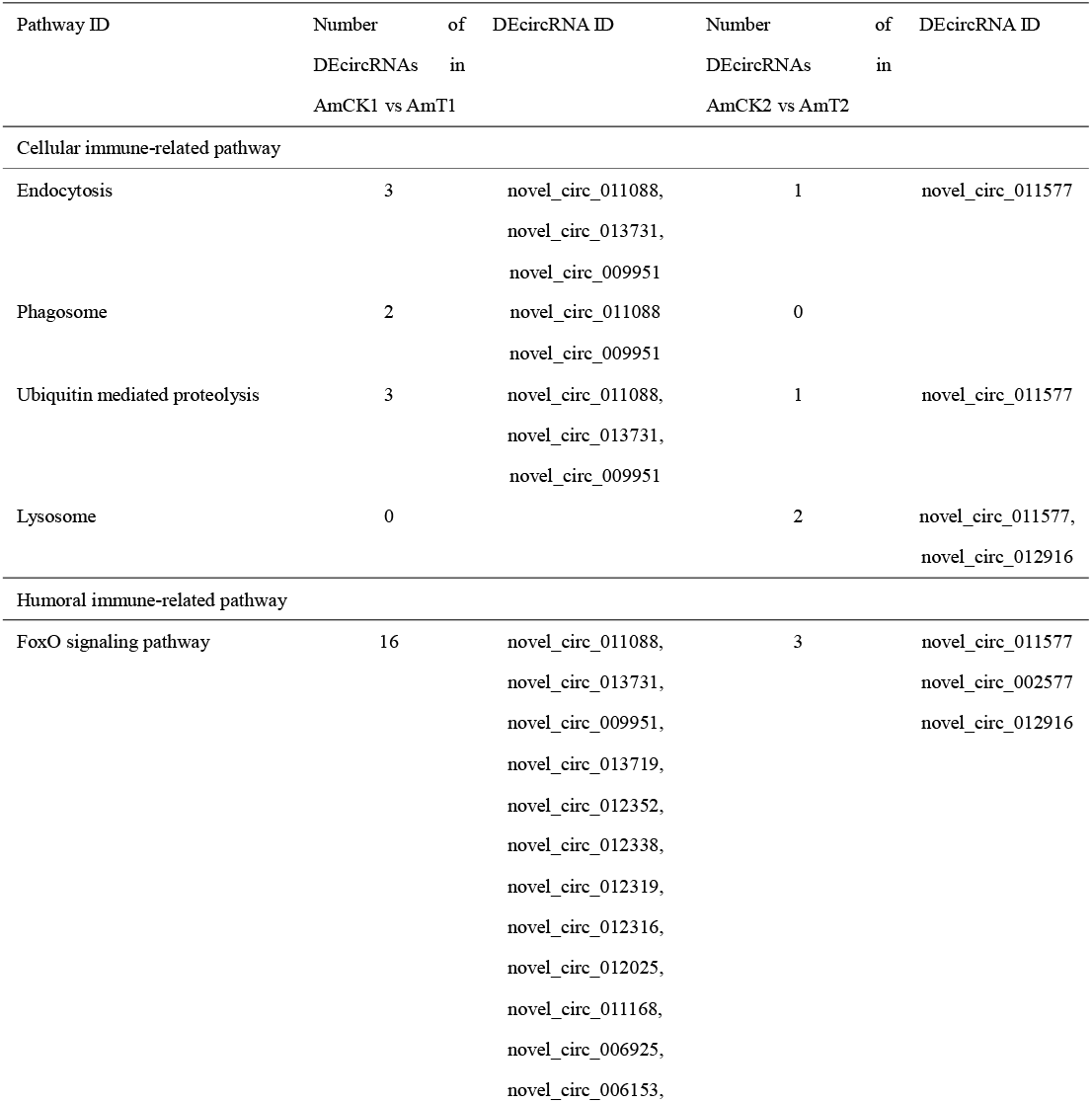
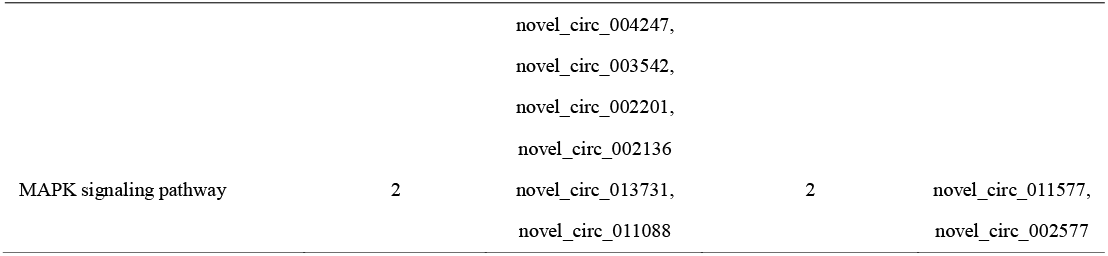
Summary of DEcircRNAs within ceRNA regulatory networks involved in host cellular and humoral immune

A model of the DEcircRNA-mediated stress response of *A. m. ligustica* workers to *N. ceranae* infection is summarized in Fig. 10. When *N. ceranae* infects the *A. m. ligustica* worker, partial DEcircRNAs in host midgut epithelial cells regulate the transcription of corresponding source genes and act as “miRNA sponges”, further participating in host metabolism, the immune response, and cell renewal such as oxidative phosphorylation, lysosome and Hippo signaling pathway.

**Fig. 10.**
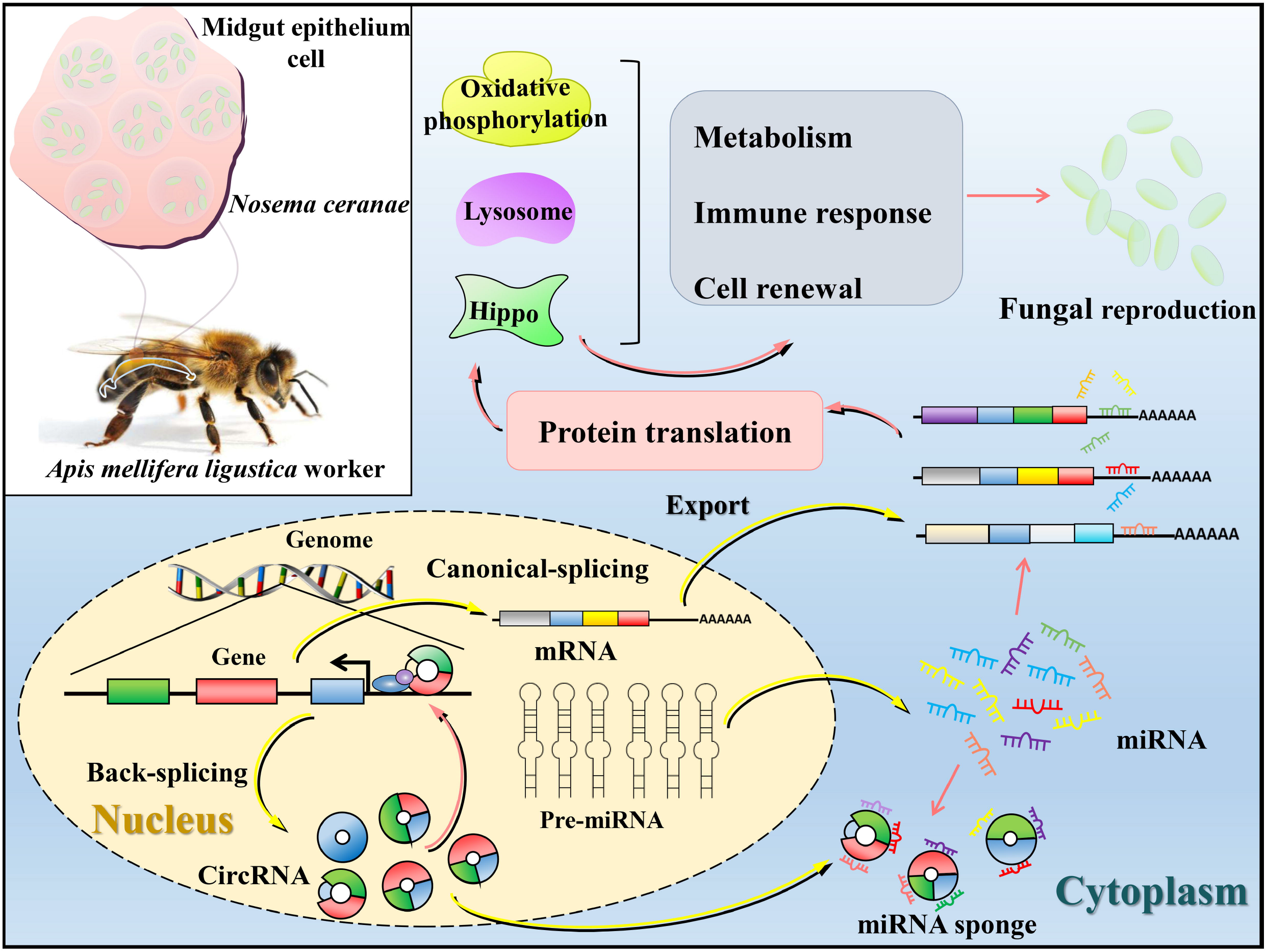
A working model showing DEcircRNA-mediated stress response of *A. m. ligustica* worker’s midgut to *N. ceranae* infection

## 4. Discussion

Currently, the mechanism regulating the immune response of western honeybees to *N. ceranae* infection is still largely unknown, although the expression of host immune genes during *N. ceranae* invasion was investigated using RT-qPCR (Antunez et al., 2009; Chaimanee et al., 2012; Li et al., 2016). To deeply explore the molecular mechanism underlying the immune response of *A. m. ligustica* workers to *N. ceranae* infection, using a collaboration of strand-specific cDNA library-based RNA-seq and sRNA-seq, we previously conducted whole transcriptome sequencing of the midguts of *A. m. ligustica* workers at 7 dpi and 10 dpi with *N. ceranae* and corresponding uninfected midguts, and on the basis of the obtained high-quality transcriptome data, we comprehensively analyzed host immune responses to *N. ceranae* infection at the mRNA, miRNA, and lncRNA levels (Fu et al., 2019; Chen et al., 2019c; Chen et al., 2020). There are two mainstream strategies to determine circRNA utilizing high-throughput sequencing, one is to construct a strand-specific cDNA library followed by deep sequencing without RNase R digestion, and the other is to digest total RNA with RNase R to remove linear RNA, followed by next-generation sequencing (Jeck and Sharpless, 2014). In the present study, the former approach was employed since the interaction between circRNAs and other RNAs such as miRNAs and lncRNAs cannot be investigated based on the linear RNA-removal strategy, although more circRNAs especially those with low expression levels, could be detected. The expression of circRNAs was specific during the developmental stage and stress response stage (He et al., 2017; Lai et al., 2018). In our previous study, 10 833 circRNAs with a length distribution between 15 nt and 1 000 nt (70.99%) were identified in the *A. m. ligustica* worker midgut using a transcriptome dataset from control groups (AmCK1 and AmCK2), and annotated-exonic circRNA was found to be the most abundant cyclization type (Xiong et al., 2018). Here, 8 199 and 8 711 circRNAs were identified from AmT1 and AmT2, respectively; additionally, 4 464 circRNAs were common to both AmT1 and AmT2, while 1 389 and 1 696 were specifically expressed in these two groups, indicative of the stress stage specificity of circRNA. Moreover, 4 464 circRNAs were found to be shared by AmCK1, AmCK2, AmT1, and AmT2. These shared circRNAs were speculated to play essential roles in not only normal midguts of *A. m. ligustica* workers but also midguts in the context of *N. ceranae* infection. At present, studies on circRNAs in western honeybees are rather limited (Guo et al., 2018b; Xiong et al., 2018; Chen et al., 2019a; Thölken et al., 2019), and the 3 409 more circRNAs identified in this study could further enrich the information on *A. mellifera* circRNAs. Combining circRNAs discovered in the control and treatment groups, a total of 14 909 *A. m. ligustica* circRNAs were finally identified, providing a valuable western honeybee circRNA reservoir for further study in the future. Further analysis suggested that 68.59% of *A. m. ligustica* circRNAs were homologous to *A. c. cerana* circRNAs, while only 0.13% had homology with human circRNAs. This indicated that the majority of circRNAs were conserved between different bee species, but the conservation of circRNAs in distant species was much lower, which was consistent with findings in other species (He et al., 2017). Interestingly, 16 circRNAs (novel_circ_002387, novel_circ_008844, novel_circ_008846, etc.) were highly conserved in all three species mentioned above, suggesting of a key function of them in both vertebrates and invertebrates, thus deserving additional investigation.

Furthermore, 168 and 306 DEcircRNAs were observed in AmCK1 vs AmT1 and AmCK2 vs AmT2, respectively, including nine common upregulated circRNAs and 10 common downregulated circRNAs; 54 (134) up-regulated circRNAs and 97 (155) down-regulated ones were specific in the aforementioned two comparison groups. This demonstrated that the expression of partial circRNAs was altered in the midgut of *A. m. ligustica* workers responding to *N. ceranae* infection, indicative of their participation in the host *N. ceranae*-response. The results suggest that the common DEcircRNAs play roles in the host midgut during fungal infection, while the unique DEcircRNAs were involved in the host response to *N. ceranae* invasion at different time points.

*N. ceranae* can destroy the structure of epithelial cells in the honeybee midgut (Doublet et al., 2015). It can also prolong the survival time of infected epithelial cells of western honeybee workers by inhibiting apoptosis, thereby exploiting material and energy from host cells for its proliferation (Panek et al., 2018; Paris et al., 2018). In the current work, 21, and two source genes in AmCK1 vs AmT1 were involved in cell renewal related-functional terms such as cellular process and cellular component organization or biogenesis; sixteen source genes (ncbi_550645, ncbi_410860, ncbi_411018, etc.) were annotated to cell structure related functional terms, such as cell component, cell, membrane, membrane part, organelle, organelle part, and membrane-enclosed lumen. In the AmCK2 vs AmT2 comparison group, more (63 and six) source genes were engaged in cellular process and cellular component organization or biogenesis; additionally, more (49) source genes (ncbi_413575, ncbi_551320, ncbi_410780 etc.) were annotated to cell part, cell, membrane, membrane part, organelle, organelle part, and membrane-enclosed lumen. Collectively, the results suggested that DEcircRNAs in the midguts of *A. m. ligustica* workers regulate genes associated with cell renewal and the cell structure of the midgut to cope with damage to the midgut epithelial cell structure caused by *N. ceranae*.

In *Drosophila*, it has been proven that Hippo signaling pathway can sense damage, deliver unpaired cytokines to stem cells, and conduct gut cell renewal (Buchon et al., 2009). In addition, the regeneration of stem cells in the *Drosophila* gut is mainly controlled by Wnt signaling pathway (Lin et al., 2008). Panek et al. found that the regeneration rate of gut stem cells of *A. mellifera* worker at 7 dpi and 14 dpi with *N. ceranae* decreased significantly owing to *N. ceranae* reproduction, and further revealed that Hippo and Wnt signaling pathways may jointly regulate stem cell differentiation in midgut epithelium (Doublet et al., 2015). In this work, we observed that two (six) and one (two) source genes of DEcircRNAs in AmCK1 vs AmT1 (AmCK2 vs AmT2) were involved in Hippo and Wnt signaling pathways, respectively. The results indicated that host DEcircRNAs might regulate the regeneration of midgut epithelial cells by regulating source genes relative to Hippo and Wnt signaling pathways, thereby responding to *N. ceranae* infection.

*N. ceranae* lacks mitochondria and hence highly depends on host ATP, which are mainly derived from the metabolism of various sugars (Chen et al., 2013). *N. ceranae* infection can increase the expression of genes related to carbohydrate metabolism in honeybees (Badaoui et al., 2017), and enhance host sensitivity of low-concentration sugar water and sugar water intake (Mayack et al., 2009). In insects, oxidative phosphorylation is the major way to synthesize ATP (Li et al., 2017). In our previous work, we reported that DElncRNAs and DEmiRNAs in the midguts of *A. m. ligustica* workers may regulate the expression levels of corresponding target mRNAs, further regulating sugar metabolism-associated pathways such as galactose metabolism and oxidative phosphorylation to respond to energy stress caused by *N. ceranae* invasion (Fu et al., 2019; Chen et al., 2019c; Chen et al., 2020). Here, two source genes of DEcircRNAs (novel_circ_013719, novel_circ_008338, novel_circ_008345, etc.) in AmCK1 vs AmT1 were engaged in galactose metabolism, including the aldose reductase coding gene (ncbi_412163) and *alpha-glucosidase* coding gene (ncbi_411257); whereas two source genes of DEcircRNAs (novel_circ_005728, novel_circ_007884, novel_circ_008164, etc.) in AmCK2 vs AmT2 were also annotated to galactose metabolism, including the *alpha glucosidase 2* coding gene (ncbi_409889) and *alpha-glucosidase* coding gene (ncbi_411257). In addition, two source genes (*alpha-glucosidase* coding gene, ncbi_411257; *trehalase*, ncbi_410484) of DEcircRNAs (novel_circ_008338, novel_circ_008345, novel_circ_012690, etc.) in AmCK1 vs AmT1 were involved in starch and sucrose metabolism, whereas four source genes of DEcircRNAs (novel_circ_010981, novel_circ_005728, novel_circ_007884, etc.) in AmCK2 vs AmT2 were also annotated to this sugar metabolism-related pathway, including the *UDP-glucuronosyltransferase 2 A3* coding gene (ncbi_409203), *alpha glucosidase 2* coding gene (ncbi_409889), *alpha-glucosidase* coding gene (ncbi_411257), and *trehalase* coding gene (ncbi_410484). Only one source gene (*aldose reductase* coding gene, ncbi_412163) of DEcircRNA (novel_circ_013719) in AmCK1 vs AmT1 was engaged in fructose and mannose metabolism. Intriguingly, only one source gene (*V-type proton ATPase 116 kDa subunit a* coding gene, ncbi_412810) of DEcircRNA (novel_circ_006925) in the host midgut at 7 dpi with *N. ceranae* was annotated to oxidative phosphorylation pathway, a key energy metabolism pathway in honeybees. Together, these results indicated that western honeybee worker may regulate sugar metabolism and oxidative phosphorylation by controlling the transcription of corresponding source genes by altering the expression of partial DEcircRNAs, thus participating in the host response with DElncRNAs and DEmiRNAs to energy stress triggered by *N. ceranae*. However, more efforts are required to clarify the underlying mechanism.

At the individual level, honeybees are able to resist pathogen invasion through cellular and humoral immunity, which is similar to other insects (Lavine et al., 2002). For honeybees, the cellular immune system is mainly composed of endocytosis, encapsulation, phagosome, melanization, and enzymatic hydrolysis, while the humoral immune system consists of synthesis and secretion of antibacterial peptides (Lavine et al., 2002). Endocytosis and phagosome are two major cellular immune pathways of honeybees (Aronstein et al., 2010). Lysosome have antifungal and antiviral activity, allowing hosts to dissolve pathogenic proteins delivered by endocytosis and phagosome (Samaranayake et al., 2001; Sardiello et al., 2001). As a significant way to clean up damaged or unwanted cells, ubiquitin-proteasome degradation system plays a primary role in cellular processes such as the cell cycle, division, differentiation, development, and immunity (McBride et al., 2003). Insect cytochrome P450 can participate in the insect immune response by catalyzing the biosynthesis and degradation of endogenous substances, including ecdysone and juvenile (Scott et al., 1998). In this research, we observed that, in the *A. m. ligustica* worker midgut at 7 dpi with *N. ceranae*, four source genes of DEcircRNAs were involved in endocytosis, including *phosphatidylinositol 4-phosphate 5-kinase type-1 alpha* coding gene (ncbi_724991), *E3 ubiquitin-protein ligase CBL* coding gene (ncbi_411982), *arrestin red cell* coding gene (ncbi_409006), and *EH domain-containing protein 3* coding gene (ncbi_413012); three source genes were engaged in phagosome, including *V-type proton ATPase 116 kDa subunit a* coding gene (ncbi_412810), *actin related protein 1* coding gene (ncbi_406122), and *ras-like GTP-binding protein RhoL* coding gene (ncbi_552419); two source genes were associated with lysosome, including *V-type proton ATPase 116 kDa subunit a* coding gene (ncbi_412810) and *battenin* coding gene (ncbi_410860). In the AmCK2 vs AmT2 comparison group, more (eight) source genes were found to be enriched in endocytosis pathway; additionally, eight (*G protein-coupled receptor kinase 1* coding gene, *ras-related protein Rab-5C* coding gene, *actin-related protein 2/3 complex subunit 1A* coding gene, etc.), two (*ras-related protein Rab-5C* coding gene, and *actin related protein 1* coding gene), one (*E3 ubiquitin-protein ligase TRIP12* coding gene), and one (*UDP-glucuronosyltransferase 2A3* coding gene) source genes of DEcircRNAs were annotated to endocytosis, phagosome, ubiquitin-mediated proteolysis, and metabolism of xenobiotics by cytochrome P450, respectively; however, no source gene was detected to be involved in lysosome. This suggested that different cellular immune pathways were regulated by modulating the transcription of various source genes by corresponding DEcircRNAs in the host midgut response to *N. ceranae* infestation. In the human FoxO signaling pathway, FoxO proteins regulate the expression of a series of target genes and participate in the immune response (Greer and Brunet, 2005). In the nucleus, the FoxO transcription factor promotes transcription of the *Bim* gene, which mediates apoptosis, leading to apoptosis (Dijkers et al., 2000). Based on our pathway analyais, we observed only one humoral immune-related pathway (FoxO signaling pathway) enriched by source genes of DEcircRNAs in the midguts of *A. m. ligustica* workers at 7 dpi and 10 dpi with *N. ceranae* infection. However, there were no source genes enriched in four classic humoral immune pathways of *A. mellifera* predicted by Evans et al. (2006), such as Toll, Imd, JAK/STAT, and JNK signaling pathways. This indicated that FoxO signaling pathway was employed by the midgut of *A. m. ligustica* workers in response to *N. ceranae* invasion via regulation of the transcription of source genes by corresponding DEcircRNAs. Together, these results suggested the participation of partial DEcircRNAs in host cellular and humoral immune responses to *N. ceranae* infection by regulating the transcription of corresponding source genes. In addition, given that target genes related to cellular immune pathways (endocytosis, ubiquitin mediated proteolysis, lysosome, etc.) and humoral pathways (FoxO signaling pathway, MAPK signaling pathway, Jak-STAT signaling pathway, etc.) were regulated by miRNAs and lncRNAs (Chen et al., 2019c; Chen et al., 2020), we speculated that western honeybees can adopt different strategies to regulate different cellular and humoral immune pathways, further responding to *N. ceranae* invasion.

Accumulating evidence has shown that circRNAs can act as “molecular sponges” to absorb miRNAs, thereby affecting the expression of downstream target genes (Huang et al., 2017; Li et al., 2018). Huang et al. (2017) discovered that circHIPK2 in mice could inhibit miR124-2HG activity via competitively targeting mir124-2HG, resulting in upregulation of the downstream *sigma non-opioid intracellular receptor 1 (SIGMAR1*) gene; knockout of circHIPK2 regulated autophagy and endoplasmic reticulum stress by altering the expression of miR124-2HG and *SIGMAR1*, further suppressing the activation of astrocytes. On the basis of high-throughput sequencing and comparative analysis of esophageal squamous cell carcinoma tissues and normal tissues, Li and colleagues found a significant upregulation of CIRS-7 in the former, and overexpression of CIRS-7 *in vitro* inhibited miR-7-mediated cell proliferation, migration, and invasion, while overexpression of CIRS-7 inhibited tumor growth and lung metabolism; the authors further revealed that CIRS-7, as a “molecular sponge” of miR-7, played a biological role by reactivating the downstream *HOXB13* gene and its NF-κB/P65 pathway (Li et al., 2018). In the past decade, researchers have made substantial progress in the honeybee miRNA field, which showing that miRNA-mediated regulation is involved in honeybee neural development (Hori et al., 2011), labor division (Liu et al., 2012), grade differentiation (Shi et al., 2015), and immune defense (Chen et al., 2020). For example, Huang et al. (2015) investigated miRNAs in *A. mellifera* workers’ midguts during six days of *N. ceranae* infection and revealed that 17 significantly differentially expressed miRNAs, including ame-miR-1, may participate in the regulation of material and energy metabolism such as oxidative phosphorylation. It has previously verified been that miR-1 is engaged in immune process of *Drosophila melanogaster* (Fullaondo et al., 2012) and *Aedes aegypti* (Shrinet et al., 2014) in response to infection by various pathogens. In the current work, we noted that ten DEcircRNAs (novel_circ_002002, novel_circ_003045, novel_circ_003465, etc.) in the AmCK2 vs AmT2 comparison group could be targeted by miR-1-z, indicating that these DEcircRNAs were likely to be involved in energy metabolism and the immune response of workers at 10 dpi with *N. ceranae* infection. Hua’s group previously found that miR-25 was correlated with the invasion and proliferation of esophageal squamous cell carcinoma (ESCC) cells; the high expression of miR-25 can result in a decreased survival rate and that the alteration of miR-25 significantly suppressed the invasion and proliferation of ESCC cells. They further found that miR-25 exerts biological function by inhibiting the expression of oncogene *F-box and WD repeat domain-containing 7 (FBXW7*) (Hua et al., 2017). Wang’s group raveled that the expression level of miR-92a in HCC cells was significantly higher than that of normal HCC cells; overexpression of miR-92a promoted the growth and invasion of HCC cells, while knockdown of miR-92a exerted an inhibitory function. Following further investigation, miR-92 was found to play its biological role via inhibition of the expression of the *Forkhead Box A2 (FOXA2*) gene (Wang et al., 2017). In the present study, five and seven DEcircRNAs in AmCK1 vs AmT1 comparison group were detected to respectively target miR-25-x and miR-92-x respectively; two miRNAs highly homologous to miR-25 and miR-92a. In another work, Xie et al. (2017) reported that miR-30a was downregulated in colorectal cancer (CRC) tissues and cell lines compared with normal rectal tissues and cells, overexpression of miR-30a *in vitro* inhibited proliferation of CRC cells and promoted apoptosis of cancer cells, further validating that miR-30a can bind to the cancer-related *ecto-5’-nucleotidase (CD73*) gene and that overexpression of *CD73* could rescue miR-30a-induced inhibition of CRC cell proliferation. Here, we noted that eight and seven DEcircRNAs in the host midgut at 7 dpi with *N. ceranae* invasion can target miR-30-x and miR-30-y, respectively; two western honeybee miRNAs shared high homology with miR-30a; whereas 12 DEcircRNAs in the host midgut at 10 dpi jointly linked to miR-30-y. Taken together, these results demonstrated that the corresponding DEcircRNAs of honeybees may be engaged in the host response to *N. ceranae* infection by absorbing miR-25-x, miR-30-y, and miR-92-x as “molecular sponges”. However, the biological function of the associated DEcircRNAs remains to be clarified.

In this work, a ceRNA regulatory network was further constructed followed by investigation of the involved target mRNAs. The results demonstrated that 86 DEcircRNAs in the host midgut at 7 dpi with *N. ceranae* infection can target 75 miRNAs, further targeting 215 mRNAs, while 178 DEcircRNAs in the host midgut at 10 dpi with *N. ceranae* infection can target 103 miRNAs, further targeting 305 mRNAs. GO database annotation showed that the abovementioned target mRNAs could be annotated to two (two) cell renewal-related terms including 50 (86) mRNAs, and seven (six) cell structure-related terms including 42 (60) target mRNAs; additionally, 12 and one target mRNAs involved in the ceRNA regulatory network in the AmCK1 vs AmT1 comparison group could be annotated to respond to stimulus and cell killing, respectively. In the AmCK2 vs AmT2 comparison group, 18 targets were also enriched in response to stimulus. Furthermore, KEGG database annotation showed that target mRNAs in the host midgut at 7 dpi with *N. ceranae* infection were engaged in 41 pathways, including two gut development-related pathways, Hippo and Wnt signaling pathways, two sugar metabolism-related pathways (amino sugar and nucleotide sugar metabolism), and one energy metabolism-related pathway such as oxidative phosphorylation pathway; while target mRNAs in the host midgut at 10 dpi were involved in two gut development-associated pathways, four sugar metabolism-associated pathways, but no energy metabolism-associated pathways. This indicated that the *A. m. ligustica* worker may systematically respond to *N. ceranae* invasion through a ceRNA mechanism by altering of partial circRNAs to regulate downstream target genes’ expression. In addition, three cellular immune pathways including endocytosis, phagosome, and ubiquitin-mediated proteolysis, as well as two humoral immune pathways, including FoxO and MAPK signaling pathways, were enriched by targets within the ceRNA network in the host midgut at 7 dpi, while endocytosis, lysosome, ubiquitin-mediated proteolysis, FoxO signaling pathway, and MAPK signaling pathway were annotated by targets within the ceRNA network within the host midgut at 10 dpi. Collectively, the results indicated the involvement of a portion of DEcircRNAs in regulating the host cellular and humoral immune response to *N. ceranae* infection by changing the expression of corresponding downstream targetgenes through the ceRNA regulatory network. Additional work is required to clarify the underlying molecular mechanism.

Apart from circRNA, several lines of evidence demonstrated that other ncRNA species, such as lncRNAs, can competitively link to miRNAs, further regulating gene expression in animals, including insects (Wang et al., 2019; Wei et al., 2019; Chen et al., 2019a; Zhang et al., 2020). Wei et al. (2019) found that miR-633a-5p can interact with nine circRNAs, eight lncRNAs, and 46 mRNAs to form a ceRNA network, while miR-154-3p interacted with five circRNAs, two lncRNAs, and 11 mRNAs to form another network, which may play a pivotal role in autophagy in pancreatic cancer cells. Chen et al. (2019a) found that 1 078 mRNAs were predicted to be targeted by 54 common miRNAs, along with 173 lncRNAs and 161 circRNAs, which were significantly enriched in Wnt signaling pathway involved in caste determination and the ovary development status in honey bees. Combined with our previous investigations of differentially expressed lncRNAs, miRNAs, and mRNAs involved in *A. m. ligustica* workers’ midgut response to *N. ceranae* infection (Fu et al., 2019; Chen et al., 2019c; Chen et al., 2020), we further constructed and analyzed cellular and humoral immune-associated DElncRNA/DEcircRNA-DEmiRNA-DEmRNA regulatory networks. We discovered that 16 DEcircRNAs and 84 DElncRNAs (data not shown) in the AmCK1 vs AmT1 comparison group could target 15 DEmiRNAs, further regulating 10 DEmRNAs relative to endocytosis, ubiquitin-mediated proteolysis, phagosome, FoxO signaling pathway, and MAPK signaling pathway, while three DEcircRNAs and 101 DElncRNAs (data not shown) in the AmCK2 vs AmT2 comparison group were found to target 26 DEmiRNAs, further regulating ten DEmRNAs relative to lysosome, ubiquitin-mediated proteolysis, endocytosis, MAPK signaling pathway, and FoxO signaling pathway. These results together unraveled complicated ceRNA regulatory networks involved in cellular and humoral immune response of western honeybee workers’ midguts to *N. ceranae* invasion, but also suggested that various ncRNAs were employed by the host to regulate immune pathways and associated genes during immune defense. The underlying mechanism deserves further exploration.

## 5. Conclusion

In the present study, for the first time, differential expression profile of circRNAs and potential regulatory role of DEcircRNAs in *A. m. ligustica* workers’ midguts response to *N. ceranae* infection especially host DEcircRNA-mediated immune defense were comprehensively analyzed. Our data revealed that the expression of host circRNAs was altered responding to *N. ceranae* challenge, a portion of DEcircRNAs may participate in host immune response to *N. ceranae* by regulating the transcription of source genes, and some other DEcircRNAs were likely to regulate host immune response as “molecular sponges” of miRNAs and further control downstream genes’ expression; additionally, complex regulatory network was formed by DEcircRNAs, DElncRNAs, DEmiRNAs, and DEmRNAs, which were jointly involved in host cellular and humoral immune response. These findings provide not only a foundation for clarifying the molecular mechanism regulating immune response of *A. m. ligustica* workers to *N. ceranae* invasion, but also a new insight into further understanding the host-pathogen interaction during bee microsporidiosis.

## Acknowledgements

This work was funded by the National Natural Science Foundation of China (31702190), the Earmarked Fund for Modern Agro-industry Technology Research System (CARS-44-KXJ7), the Science and Technology Planning Project of Fujian Province (2018J05042), the Education and Scientific Research Program Fujian Ministry of Education for Young Teachers (JAT170158), the Outstanding Scientific Research Manpower Fund of Fujian Agriculture and Forestry University (xjq201814), the Master Supervisor Team Fund of Fujian Agriculture and Forestry University, and the Fund for Excellent Master Dissertation of Fujian Agriculture and Forestry University. RG genuinely appreciates and cherishes the memory of his beloved father.

---

**Table S1** Divergent primers and convergent primers PCR validation of novel circRNAs and DEcircRNA

**Table S2** Primers for Stem-loop PCR validation of circRNA target miRNAs

**Table S3** DEcircRNAs in AmCK1 vs AmT1 comparison group

**Table S4** DEcircRNAs in AmCK2 vs AmT2 comparison group

**Table S5** GO database annotation of DEcircRNA source genes in AmCK1 vs AmT1 comparison group

**Table S6** GO database annotation of DEcircRNA source genes in AmCK2 vs AmT2 comparison group

**Table S7** KEGG database annotation of DEcircRNA source genes in AmCK1 vs AmT1 comparison group

**Table S8** KEGG database annotation of DEcircRNA source genes in AmCK2 vs AmT2 comparison group

**Table S9** Relationship between DEcircRNAs and miRNAs in AmCK1 vs AmT1 comparison group

**Table S10** Relationship between DEcircRNAs and miRNAs in AmCK2 vs AmT2 comparison group

**Table S11** Relationship between miRNA targeted by DEcircRNA and mRNA in AmCK1 vs AmT1 comparison group

**Table S12** Relationship between miRNA targeted by DEcircRNA and mRNA in AmCK2 vs AmT2 comparison group

**Table S13** GO database annotations of target mRNAs in AmCK1 vs AmT1 comparison group

**Table S14** GO database annotations of target mRNAs in AmCK2 vs AmT2 comparison group

**Table S15** KEGG database annotations of target mRNAs in AmCK1 vs AmT1 comparison group

**Table S16** KEGG database annotations of target mRNAs in AmCK2 vs AmT2 comparison group

**Fig. S1. Pearson correlation coefficients between different biological replicates in AmT1 (A) and AmT2 (B).**

