## Supplementary figures and images for "Deciphering the mechanism underlying circRNA-mediated immune responses of western honeybees to *Nosema ceranae* infection"

### Fig S1

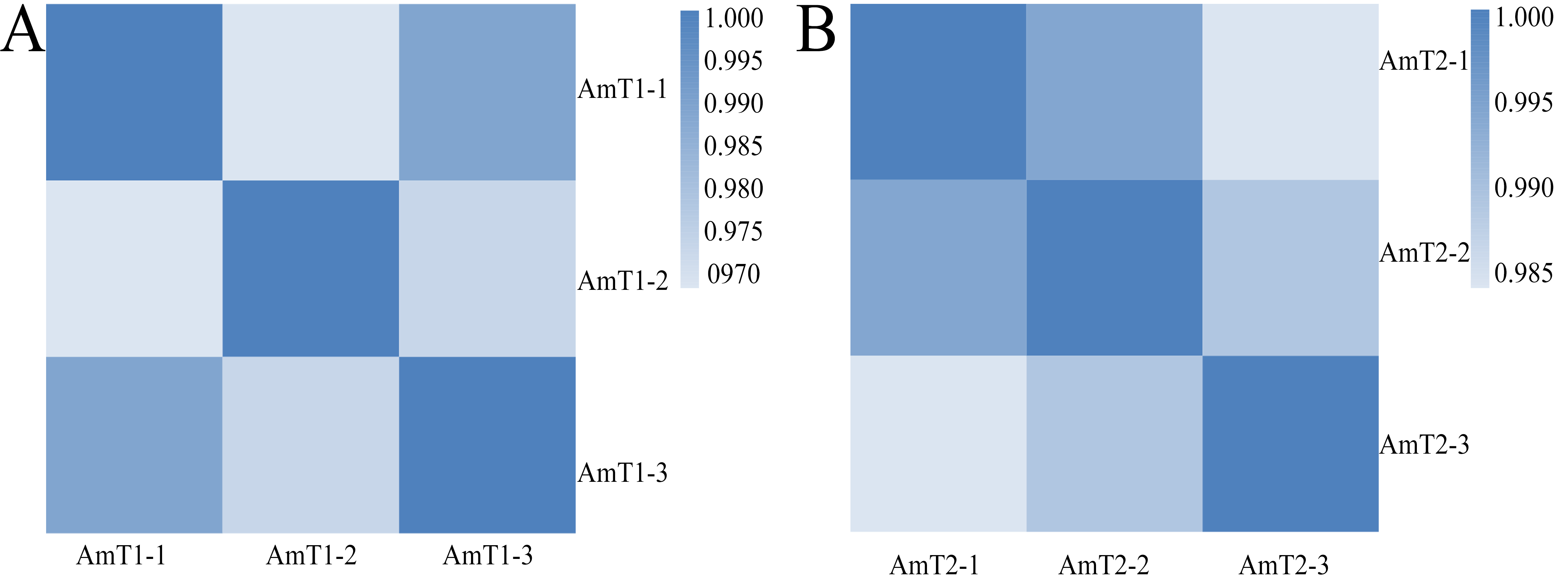
