## Supplementary material for "Deciphering the mechanism underlying circRNA-mediated immune responses of western honeybees to *Nosema ceranae* infection": Table S1

**Table S1** Divergent primers and convergent primer of molecular validation of novel circRNAs and DEcircRNAs

| Id | Type of primers | Sequence | Product size (bp) | Purpose |
| --- | --- | --- | --- | --- |
| Novel_circ_004065 | Divergent Primer | F: CTCCACAATGTTTACCTGGTC | 86 | RT-PCR |
|  |  | R: ATCTCCGAGTTCTTGAGCG |  |  |
|  | Convergent primer | F: TCCTCATTCCCTCATCAATC | 301 |  |
|  |  | R: CGCTTTCAGCAGATACTCG |  |  |
| Novel_circ_002199 | Divergent Primer | F: AACAGCGTTGAATCAGGC | 187 | RT-PCR |
|  |  | R: TTCGGGCAAAGGATGTAG |  |  |
|  | Convergent primer | F: GGAAGAAGAAGCGAGCAAG | 358 |  |
|  |  | R: TACAATGGGAGAGTCAGTGG |  |  |
| Novel_circ_005784 | Divergent Primer | F: ACACCTGCTGCTACACCATC | 91 | RT-PCR |
|  |  | R:GCTGGAGTTACTGCGTATCC |  |  |
|  | Convergent primer | F: CATACGATTACGGTTACGGAC | 312 |  |
|  |  | R: TTGTCGCTGTTGATGGTG |  |  |
| Novel_circ_000705 | Divergent Primer | F: CAAGTCCAAGGCGAAGAAG | 259 | RT-qPCR |
|  |  | R: CCCTCGGTGTTCAACTGTAG |  |  |
|  | Convergent primer | F: ATCTTCTGGACACCATCAGC | 264 |  |
|  |  | R: TCTCGTCTTTGGATTCTCG |  |  |
| Novel_circ_001195 | Divergent Primer | F: ATAATGGCGGTTCGCTGAG | 220 | RT-qPCR |
|  |  | R: ATTGCCTAACAAAGTTGGAGGG |  |  |
|  | Convergent primer | F: TCTCGCATTGTTCTCAGGG | 351 |  |
|  |  | R: AAGCGGTTTCCTCTCGTCAC |  |  |
| Novel_circ_011173 | Divergent Primer | F: AGAGCGTGGAAAGCAGAAC | 255 | RT-qPCR |
|  |  | R: GGAAAGAGAAAGAATGGTCG |  |  |
|  | Convergent primer | F: GGAGAAAGAAAGAATGCGTG | 455 |  |
|  |  | R: GCCTGCTTGAAGATTTGC |  |  |
| Novel_circ_006925 | Divergent Primer | F: CCCTTATCCGTTGGGTATG | 244 | RT-qPCR |
|  |  | R: ACAAAGAGGCGTGGAAAC |  |  |
|  | Convergent primer | F: TCCACGCCTCTTTGTATCC | 290 |  |
|  |  | R:GCAGTCTCTGACGATGGTAAG |  |  |
| Novel_circ_012352 | Divergent Primer | F: GCTACCGTATTGCCATTCAC | 133 | RT-qPCR |
|  |  | R: ACATTGATGCTGGTGTCG |  |  |
|  | Convergent primer | F: ACGAAACGACGACGAGTTC | 344 |  |
|  |  | R: TGAATGGCAATACGGTAGC |  |  |
|  |  | R: GTGATTGCTGTTGTCGTTG |  |  |
| Novel_circ_012316 | Divergent Primer | F: CCTGTCTCTCCACAAATGTTTC | 244 | RT-qPCR |
|  |  | R: TGTTCTCACGGTTACGGAC |  |  |
|  | Convergent primer | F: CTATGCCACCCATTCCTAAC | 324 |  |
|  |  | R: ATGTAAACGGCGGTCTGAC |  |  |
| Novel_circ_007686 | Divergent Primer | F: AACAATACACTTGCCCAGG | 131 | RT-qPCR |
|  |  | R: TCTTCCGCCAATCTGAAC |  |  |
|  | Convergent primer | F: TCGGTTTCGTTAGGGAGAC | 312 |  |
|  |  | R: TGGACACCAGATGATTCG |  |  |
| Novel_circ_011500 | Divergent Primer | F: GCAATCCAAGGACAATCTG | 166 | RT-qPCR |
|  |  | R: TATTCCAGTCTGTGGGCTG |  |  |
|  | Convergent primer | F: AACAGCATCAGCAACAAGC | 130 |  |
|  |  | R: GGATTTCCGTTCCTAAGTGC |  |  |
| *actin* |  | F: CACTCCTGCTATGTATGTCGC | 132 | RT-qPCR |
|  |  | R: GGCAAAGCGTATCCTTCA |  |  |
