## Supplementary material for "Deciphering the mechanism underlying circRNA-mediated immune responses of western honeybees to *Nosema ceranae* infection": Table S2

**Table S2** Primers for Stem-loop PCR validation of circRNA target miRNAs

| Primer name | Primer sequence (5’-3’) |
| --- | --- |
| Ame-mir-3720-loop | CTCAACTGGTGTCGTGGAGTCGGCAATTCAGTTGAGCTGTTTAA |
| Mir-21-x-loop | CTCAACTGGTGTCGTGGAGTCGGCAATTCAGTTGAGGTCAACAT |
| Mir-30-x-loop | CTCAACTGGTGTCGTGGAGTCGGCAATTCAGTTGAGAGCTTCCA |
| Mir-29-y-loop | CTCAACTGGTGTCGTGGAGTCGGCAATTCAGTTGAGTAACCGAT |
| Mir-451-x-loop | CTCAACTGGTGTCGTGGAGTCGGCAATTCAGTTGAGACTCAGTA |
| Mir-7975-y-loop | CTCAACTGGTGTCGTGGAGTCGGCAATTCAGTTGAGTGGTGCCG |
| Mir-146-x-loop | CTCAACTGGTGTCGTGGAGTCGGCAATTCAGTTGAGCCATCTAT |
| Mir-143-y-loop | CTCAACTGGTGTCGTGGAGTCGGCAATTCAGTTGAGGAGCTACA |
| Mir-101-y-loop | CTCAACTGGTGTCGTGGAGTCGGCAATTCAGTTGAGCTTCAGTT |
| Mir-462-x-loop | CTCAACTGGTGTCGTGGAGTCGGCAATTCAGTTGAGCAGCTGCA |
| Ame-mir-3720-F | ACACTCCAGCTGGGATACGGTGATGAGT |
| Mir-21-x-F | ACACTCCAGCTGGGTAGCTTATCAGACTG |
| Mir-30-x-F | ACACTCCAGCTGGGTGTAAACATCCTCGAC |
| Mir-29-y-F | ACACTCCAGCTGGGTAGCACCATCTGAA |
| Mir-451-x-F | ACACTCCAGCTGGGAAACCGTTACCAT |
| Mir-7975-y-F | ACACTCCAGCTGGGATCCTGGTCA |
| Mir-146-x-F | ACACTCCAGCTGGGTGAGAACTGAATTCC |
| Mir-143-y-F | ACACTCCAGCTGGGTGAGATGAAGCAC |
| Mir-101-y-F | ACACTCCAGCTGGGTACAGTACTGTGAT |
| Mir-462-x-F | ACACTCCAGCTGGGTAACGGAACCCATAA |
| Universal R | CTCAACTGGTGTCGTGGA |
